# Lipid Headgroup Hydration Regulates Distinct Remodeling of the Membrane Interface by Polyethylene Glycol and Dextran

**DOI:** 10.64898/2026.07.18.739366

**Authors:** Sayani Bhunia, Jacky Lin, Dmytro Nykypanchuk, Simou Sun

**Affiliations:** Department of Chemistry, Stony Brook University, Stony Brook, NY 11733, United States; Center for Functional Nanomaterials, Brookhaven National Laboratory, Upton, New York 11973, United States

## Abstract

Water-soluble polymers commonly interact with cell membranes, but their interactions are poorly understood. Here, we investigate polyethylene glycol (PEG) and dextran (DEX) interactions with different model lipid membranes. Using total internal reflection fluorescence microscopy, we observe that PEG and DEX trigger strikingly different membrane responses – DEX induces extensive membrane remodeling, including localized multilamellar domain formation, while PEG does not. Combining fluorescence spectroscopy, fluorescence anisotropy, and vibrational sum frequency spectroscopy, we show that DEX perturbs lipid headgroup hydration by displacing interfacial water with minimal effects on lipid packing, while PEG largely preserves this hydration layer. We find that membrane binding affinity alone does not determine the extent to which hydrophilic polymers perturb membrane structure and interfacial properties; and that lipid headgroup hydration, rather than lipid charge, is a general regulator of hydrophilic polymer-membrane interactions. This work gives mechanistic insights into how neutral polymers interact with cells and vesicles, with relevance to cell biology and drug delivery.

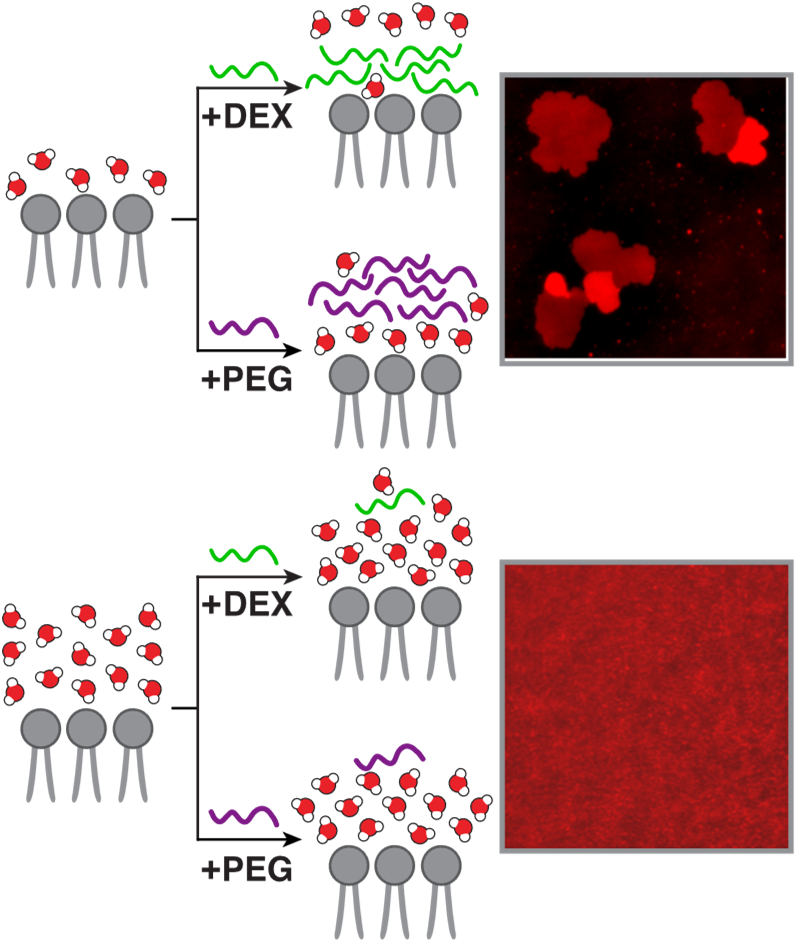

## Introduction

Water-soluble polymers ubiquitously interface with biomembranes and play critical roles in regulating membrane-associated biological processes. Examples include glycans that form a surface coating on virtually all cells,^1^ polymer-based biomaterials that function through direct contact with biological systems,^2, 3^ and polymer-functionalized drug delivery vehicles.^4^ Despite extensive investigations of their physicochemical properties and biological functions, most previous studies have focused on polymer– protein interactions and have been conducted either in the absence of lipid membranes or by treating membranes as passive substrates. Increasing evidence, however, indicates that many natural and bio-inspired polymers generate crowding that can drive lipid vesicle clustering and membrane fusion in the bulk solution,^8–11^ as well as membrane deformation on a two-dimensional surface.^12, 13^ Moreover, some polymers can directly interact with lipid membranes^5–7^. Nevertheless, the molecular principles that govern polymer–membrane interactions remain poorly understood, particularly how membrane physicochemical properties actively regulate polymer adsorption, and the functional consequences for local interface environment have yet to be elucidated. This knowledge gap limits our ability to understand and predict how water-soluble polymers control membrane-associated biological processes and to rationally engineer polymer-based biomaterials with programmable functions.

Most existing studies of lipid–polymer interactions have focused on how the physical and chemical properties of the polymers govern these interactions.^14, 15^ Consequently, the influence of membrane properties and lipid composition has received considerably less attention. Moreover, previous work has primarily examined polymers with complex chemistries, such as branched architectures, charged polymers, and block copolymers, which often exhibit relatively strong membrane interactions.^6, 7, 16^ In contrast, a broad class of hydrophilic polymers with simpler chemistries, including uncharged and predominantly linear polymers, is generally assumed to interact only weakly with lipid membranes.^6, 17^. However, several intriguing recent observations suggest that these seemingly weak interactions are tunable and may have important biological consequences. For example, one study demonstrated that increasing the hydrogen-bonding capacity of membrane lipids enhances their interactions with hydrophilic polymers.^18^ Another study showed that a pre-coated glycan layer can modulate lipid phase separation in membranes formed atop the glycan network: heterogeneous glycan networks stabilize large lipid domains at the network length scale, whereas homogeneous networks suppress macroscopic domain formation.^19^ Although the molecular mechanism underlying this glycan-patterning effect remains unclear, a more recent study reported that a homogeneous hyaluronic acid layer preferentially associates with domain-forming lipids and promotes the formation of sphingomyelin- and cholesterol-rich domains.^20^ Collectively, these findings suggest that the physical and molecular principles governing weak but direct lipid-polymer interactions - and how these interactions depend on membrane composition - remain poorly understood.

In this work, we characterize the interactions of two uncharged, hydrophilic polymers, polyethylene glycol (PEG) and dextran (DEX), with lipid membranes and investigate how lipid headgroup chemistry regulates these interactions. PEG and DEX are widely used in biomaterials that interface directly with biological systems, serving as common molecular crowding agents in biochemical assays and as surface-functionalization components for drug delivery vehicles.^21, 22^ Interestingly, the two polymers are fully miscible at low concentrations, forming a homogeneous aqueous solution. However, above critical concentrations, they undergo phase separation into PEG-rich and DEX-rich aqueous phases, giving rise to the classical aqueous two-phase system (ATPS).^23^ Consequently, PEG/DEX mixtures have been widely used to construct lipid-based artificial cells and protocell models and to investigate questions including the origin of life.^24–26^ Despite their widespread use, both polymers are generally considered to interact only weakly with zwitterionic phosphatidylcholine (PC) membranes under ambient conditions. For example, the binding free energy of PEG was reported to be 1.6 k_B_T^27^ with an enthalpy change below the detection limit of ITC.^6^ Moreover, PEG has consistently been reported to exhibit a greater affinity towards PC membranes than DEX.^28–30^

We first directly visualize and quantify the interactions of PEG or DEX with model lipid membranes using fluorescently labeled polymers and total internal reflection fluorescence microscopy (TIRFM) on supported lipid bilayers (SLBs). We show that lipid headgroup chemistry strongly regulates the binding of both polymers: incorporation of the weakly hydrated cationic lipid 1,2-dioleoyl-3-trimethylammonium-propane (DOTAP) enhances polymer adsorption, whereas the strongly hydrated anionic lipid 1,2-dioleoyl-*sn*- glycero-3-phospho-(1′-*rac*-glycerol) (DOPG) slightly suppresses it. Despite exhibiting similar trends in membrane binding, PEG and DEX produce markedly different effects on membrane organization, with only DEX inducing extensive membrane remodeling, including lamellar-to-multilamellar structural transitions. Combining fluorescence spectroscopy, fluorescence anisotropy, and vibrational sum frequency spectroscopy (VSFS), we further demonstrate that DEX, but not PEG, substantially perturbs the hydration environment of DOTAP headgroups, whereas neither polymer notably alters the hydration of DOPG-containing membranes or membrane lipid packing. Extending these observations to additional lipid systems, we identify lipid headgroup hydration as a general molecular regulator controlling direct interactions between hydrophilic polymers and lipid membranes. Finally, temperature-dependent binding measurements provide thermodynamic insight into the distinct membrane interactions of PEG and DEX. Together, this work establishes a molecular framework for understanding how lipid headgroup chemistry regulates polymer–membrane interactions and demonstrates that polymers with similarly weak membrane affinities can produce fundamentally different effects on membrane organization and interfacial hydration. These findings provide new insight into polymer - membrane interfaces relevant to glycocalyx biology and biological activity of biomaterials.

## Results

To directly characterize polymer-membrane interactions, we separately introduced FITC-labeled DEX (FITC-DEX, 10 kDa) and rhodamine B-labeled PEG (RhoB-PEG, 8 kDa) to SLBs, a widely used model membrane platform.^31, 32^ SLBs allow systematic tuning of lipid composition and are compatible with TIRFM, which is a microscopy imaging technique that enables selective visualization of membrane-associated fluorescent polymers (Figure 1A). PEG 8 kDa and DEX 10 kDa constitute one of the most commonly used polymer pairs in aqueous ATPS studies,^33^ due to their similar hydrodynamic radii, viscosity, and comparable osmotic activity.^10^ Therefore, we use them for this study and refer to them hereafter simply as PEG and DEX. In later sections of the paper, we also demonstrate with PEG8 kDa and PEG 10kDa exert similar impacts on the membrane (Figure S7, S16). FITC-DEX and RhoB-PEG have been widely employed as fluorescent analogues of their native polymers, particularly in studies of polymer–biomembrane interfaces.^24–26, 28, 29^ (We have also provided detailed chemical information regarding FITC-DEX and RhoB-PEG in the SI, and the justification of choosing these two dyes for DEX and PEG respectively). To evaluate whether the fluorescent labeling substantially alters physicochemical properties of the two polymers, we compared the labeled and native polymers using dynamic light scattering and zeta potential measurements. The labeled polymers exhibited similar hydrodynamic radii and near-neutral zeta potentials (Table S1), suggesting that fluorescent labeling minimally perturbs the polymers’ solution properties. Therefore, the adsorption behavior observed here is expected to largely reflect that of the native polymers, although we cannot completely exclude subtle effects arising from fluorescent labeling.

**Figure 1.**
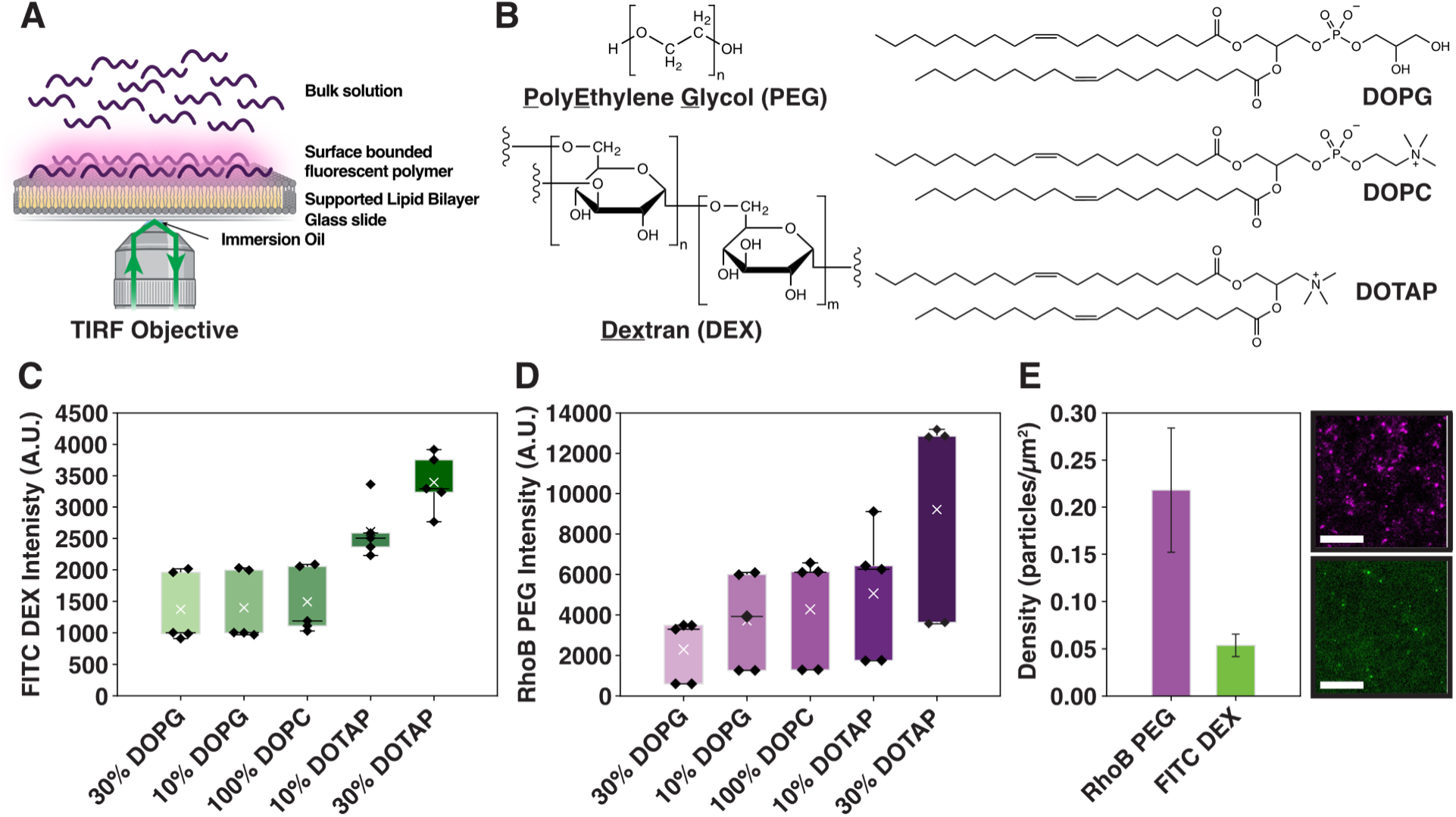
DOTAP prominently enhances polymer-membrane binding, while DOPG slightly suppresses it. (A) Schematic of the TIRF experimental setup. (B) Chemical structures of polymers and lipids used in this study. (C) TIRF intensity of surface-bound FITC–DEX (0.1 μM) as a function of membrane composition. (D) TIRF intensity of surface-bound RhoB–PEG (0.1 μM) as a function of membrane composition. Error bars represent standard deviations from five independent experiments performed on five SLBs. For DOPG/DOTAP containing SLBs, the remaining lipid composition is DOPC. (E) Left: Comparison of the surface densities of FITC–DEX and RhoB–PEG (both 10 nM) on SLBs made up of 50 mol% DOPC+ 50 mol% DOTAP (FITC–DEX particle densities were corrected for fluorophore labeling efficiency). Error bars represent standard deviations from three independent experiments performed on three SLBs. Right: Representative single-molecule TIRF images of RhoB–PEG (top) and FITC–DEX (bottom) adsorbed on the same membrane composition. Scale bars: 10 μm.

At a bulk polymer concentration of 0.1 μM, both FITC-DEX and RhoB-PEG readily adsorbed onto SLBs composed of 100 mol% zwitterionic DOPC (Figures 1B, 1C, and S2). Previous surface tension measurements reported detectable adsorption of both polymers to phosphatidylcholine membranes only at polymer concentrations above 3.33 mM.^30^ Our results therefore demonstrate that TIRFM enables detection of membrane-associated polymers at concentrations several orders of magnitude lower than those required in earlier measurements. We next varied the lipid composition by incorporating either DOTAP or DOPG into the SLBs (Figure 1B). For both polymers, adsorption was markedly enhanced by DOTAP but slightly reduced by DOPG (Figures 1C and 1D). The same lipid composition-dependent trend was observed at near-single-molecule polymer concentrations (Figure S3), indicating that the observed dependence on membrane composition is unlikely to arise from polymer–polymer interactions.

DOTAP possesses a cationic trimethylammonium headgroup and, because it lacks a phosphate group, exhibits weak hydrogen-bonding capability.^34, 35^ In contrast, DOPG contains a phosphate group and two hydroxyl groups, giving rise to an anionic headgroup with strong hydrogen-bonding capability.^34, 35^ It is therefore counterintuitive that two uncharged, hydrophilic polymers with known hydrogen-bonding capabilities adsorb more strongly to DOTAP-containing membranes than to DOPG-containing membranes.

To examine whether electrostatic interactions contribute to this behavior, we repeated the experiments in the presence of 150 mM NaCl. A similar adsorption trend was observed (Figure S4), indicating that lipid charge is unlikely to be the primary determinant of the differential polymer adsorption. The origin of this unexpected lipid dependence will be investigated in the following sections. Notably, at the same near-single-molecule polymer concentration (Figure S1), the surface density of RhoB-PEG was approximately four to five times higher than that of FITC-DEX (Figure 1E). Control experiments using free rhodamine B confirmed that the fluorophore itself does not promote membrane association (Figure S5). Furthermore, competitive binding experiments demonstrated that PEG exhibits a higher membrane affinity than DEX (Figure S6), in agreement with previous reports showing preferential partitioning of RhoB-PEG over FITC-DEX to membrane surfaces when both polymers are present simultaneously.^28–30^ Together, these results demonstrate that PEG and DEX exhibit similar lipid composition-dependent binding behavior, and PEG consistently adsorbs more strongly than DEX.

Next, we examined membrane structural changes upon addition of unlabeled PEG or DEX. SLBs consisting of 99.9 mol% DOPC and 0.1 mol% RhoB-POPE were prepared, with RhoB-POPE serving as a fluorescent membrane probe for TIRFM. Prior to polymer addition, the SLBs were laterally homogeneous, and incubation with 0.3 g/mL PEG produced no detectable changes in membrane morphology (Figure 2A). In contrast, addition of the same concentration of DEX, followed by a 5 min incubation, induced the formation of bright, micron-sized domains that continued to grow over the subsequent 1–2 h (Figure 2A and Movie S1). Control experiments with PEG 10 kDa showed no detectable patch formation (Figure S7), demonstrating the difference between DEX10K and PEG8K is not due to their different molecular weight Line profile analysis revealed discrete increases in fluorescence intensity within these domains relative to the surrounding membrane. For example, the representative line profile in Figure 2B shows that the fluorescence intensity of the bright domains is approximately 2–3-fold higher than that of the adjacent bilayer, while remaining approximately uniform across each domain (Figure 2B). Such discrete intensity increases suggest a structure of stacked lipid bilayers and are later confirmed by AFM measurements. In fact, they are reminiscent of multilamellar structures reported for SLBs under bulk dehydration conditions.^36^ Furthermore, DEX-induced membrane remodeling was strongly dependent on membrane composition. DEX promotes the formation of a high density of micron-sized domains on DOTAP-containing membranes, whereas it failed generate comparable domains on DOPG-containing membranes, aside from occasional small puncta (Figure S8).

**Figure 2.**
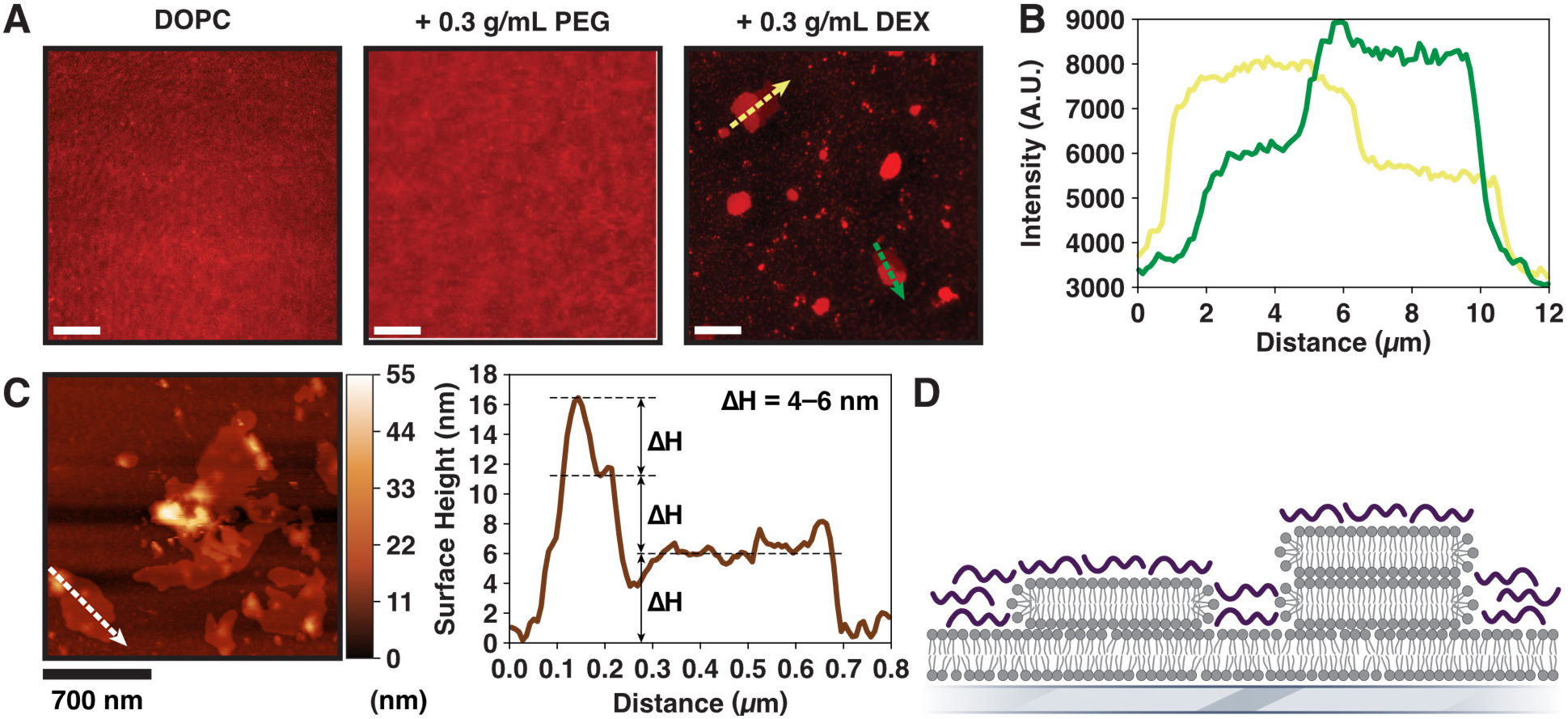
Dextran, not PEG, induces membrane stacking on SLBs. (A) TIRF images of a SLB composed of 99.9 mol% DOPC and 0.1 mol% RhoB-POPE before polymer addition (left), after addition of PEG (0.3 g/mL, center), and after addition of Dextran (0.3 g/mL, right). (B) Representative fluorescence intensity line scans taken across two membrane stacks (along the green and yellow lines). (C) Representative AFM image of DEX-induced membrane stacks formed on the same membrane composition (left) with the corresponding height profile taken along the dashed line (right). (d) Schematic illustration of the proposed model for dextran promoting local membrane stacking on the SLB. Scale bars: 10 μm (A) and 700nm (C).

To directly characterize the three-dimensional structure of these domains, we performed atomic force microscopy (AFM). AFM topographic images revealed well-defined raised domains on the membrane surface (Figure 2C, left). Height profiles acquired across these domains exhibited discrete step heights, with the surrounding SLB serving as the reference (0 nm) and additional steps of approximately 4–6 nm, 12 nm and 16 nm (Figure 2C, right). Control experiment measured the structure and height of the surrounding unilamellar SLB by creating an artificial scratch mark to expose the glass surface (Figure S9). The smallest step height (4–6 nm) is consistent with the thickness of a single phospholipid bilayer^37^ within experimental variation, while the larger step heights correspond to approximately two and three additional bilayers, respectively, indicating the formation of locally stacked multilamellar membrane domains. A schematic illustrating the proposed multilayer architecture is shown in Figure 2D. Similar membrane remodeling was observed using the tail-labeled fluorophore BODIPY-PC, demonstrating that the phenomenon is independent of both the chemical identity and labeling position of the fluorescent probe (Figure S10). Collectively, these results demonstrate that DEX, but not PEG, promotes localized membrane stacking and multilamellar domain formation under the conditions examined. Furthermore, this structural remodeling is strongly dependent on membrane composition, with DOTAP facilitating and DOPG suppressing multilayer formation, in line with the polymer adsorption trends shown in Figure 1. To investigate the molecular basis for these distinct polymer-induced membrane responses, we next examined the membrane–polymer interface using solvatochromic dye-based fluorescence spectroscopy.

Laurdan is a widely used fluorescent membrane probe that localizes to the membrane–water interface, with its dimethylamino group positioned near the glycerol carbonyl region of phospholipids (Figure 3A).^38, 39^ Its emission spectrum exhibits two broad maxima at approximately 440 and 490 nm, which are sensitive to the physicochemical properties of the membrane interface, particularly lipid packing and interfacial hydration. Reduced interfacial hydration and increased lipid packing can both increase the emission intensity at 440 nm while decreasing that at 490 nm (see detailed discussion about this mechanism in the SI).^40, 41^ These spectral changes are quantified using the generalized polarization (GP) parameter (Figure 3B), where an increase in GP corresponds to reduced interfacial hydration and/or tighter lipid packing, and consequently lower membrane fluidity. Although membrane fluidity is often used as a general descriptor of these membrane properties^38, 39^, Laurdan GP does not directly measure fluidity. Rather, it reports changes in the local physicochemical environment surrounding the probe arising from variations in interfacial hydration and lipid packing.

**Figure 3.**
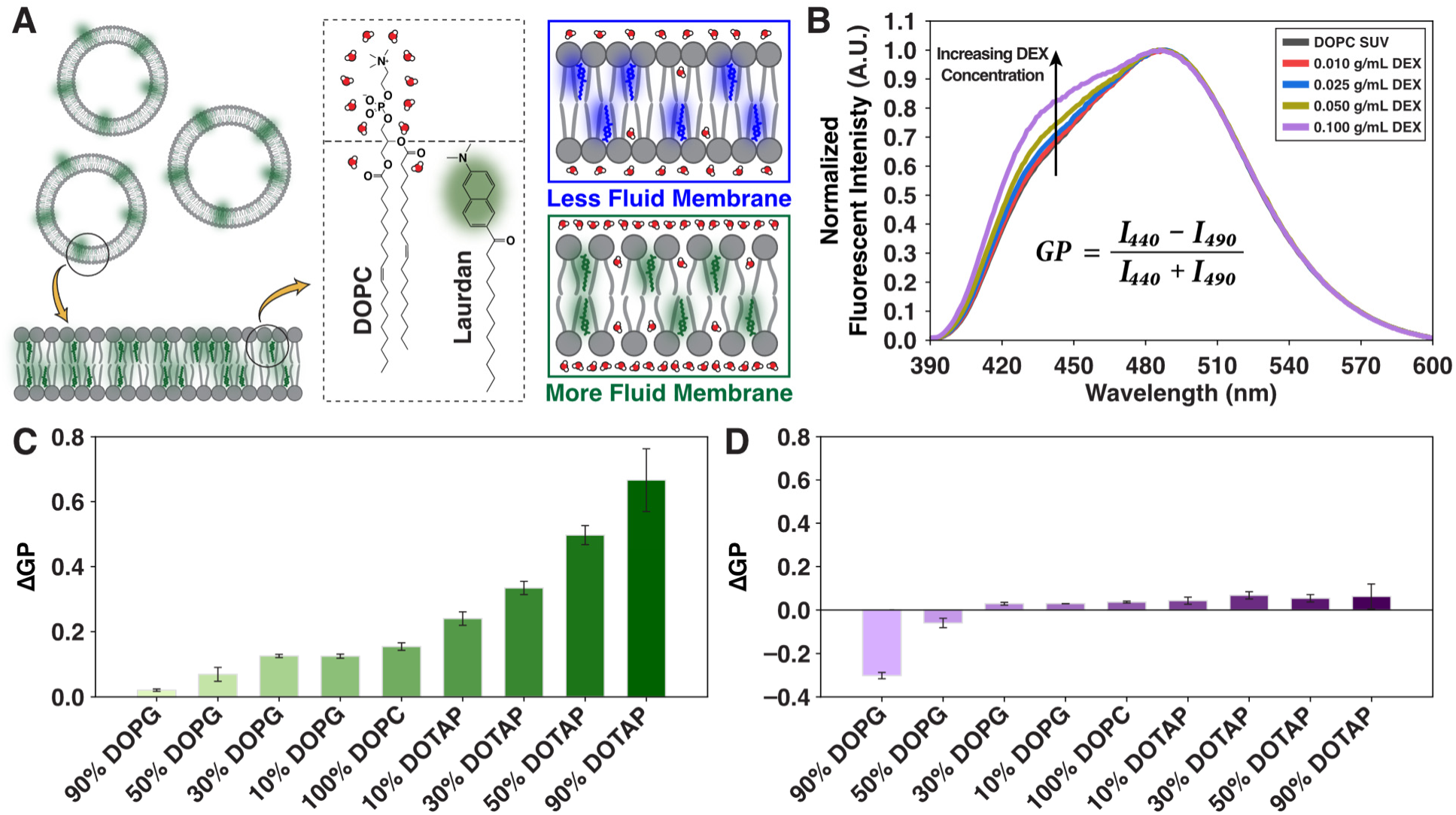
Dextran reduces membrane fluidity, while PEG shows minimal effect. (A) (Left) Schematic of the fluorometry experiment using Laurdan incorporated into SUVs. The inset shows the molecular structures of Laurdan and DOPC, highlighting the location of Laurdan within the lipid bilayer. (Right) Illustration of how Laurdan reports on membrane state: reduced interfacial hydration and/or increased lipid packing leads to a less fluid membrane. (B) Normalized emission spectra of Laurdan in DOPC SUVs with increasing dextran concentration. A generalized polarization (GP) value can be calculated from the fluorescence intensity at 440 nm and 490 nm. (C) Polymer-induced change in GP (ΔGP) as a function of membrane composition in the presence of DEX. (D) ΔGP as a function of membrane composition in the presence of PEG. For DOPG/DOTAP containing SUVs, the remaining lipid composition is DOPC. Error bars represent standard deviations from five independent experiments performed using five separate SUV preparations.

To evaluate polymer-induced changes in membrane fluidity across a broad range of membrane compositions, we incorporated Laurdan into small unilamellar vesicles (SUVs) rather than SLBs, thereby avoiding the inefficient vesicle fusion observed for membranes containing high fractions of DOPG or DOTAP. Representative Laurdan emission spectra from 100 mol% DOPC SUVs excited at 360 nm are shown in Figure 3B. Increasing the DEX concentration from 0.01 to 0.1 g/mL was introduced to the SUVs, and it progressively increased the 440 nm emission, indicating a decrease in membrane fluidity. Because water-soluble polymers such as PEG and DEX reversibly cluster vesicles through crowding-induced depletion interactions,^10^ we first ask whether polymer crowding alone could account for the observed GP changes. Consistent with previous reports, both polymers increased the apparent hydrodynamic diameter of SUVs, with PEG producing the larger effect; importantly, these changes were fully reversible upon dilution (Figure S11). Surprisingly, despite inducing substantially stronger depletion interactions, PEG caused only minimal GP increases relative to DEX (Figure 3D), indicating that the spectral changes observed in Figure 3B cannot be explained solely by polymer crowding. Because Laurdan primarily reports the hydration environment immediately surrounding the lipid carbonyl region,^38, 39^ depletion of bulk water or more loosely associated interfacial water by polymer crowding^42, 43^ is expected to have only a limited influence on its fluorescence. Moreover, SUVs with preloaded DEX produced comparable GP changes following external DEX addition (Figure S12), indicating that osmotic pressure differences across the membrane are not responsible for the observed spectral changes. Collectively, these results indicate that the observed GP changes primarily arise from alterations in the local membrane environment caused by polymer–membrane interactions rather than nonspecific crowding effects or osmotic pressure imbalances across the membrane.

We next examined how DEX alters membrane fluidity as a function of lipid composition. As shown in Figure 3C, DEX increased the GP value (ΔGP > 0) for all membrane compositions examined, indicating an overall reduction in membrane fluidity. Notably, the magnitude of ΔGP increased monotonically with increasing DOTAP content and decreased with increasing DOPG content. Thus, DEX substantially reduced the fluidity of DOTAP-rich membranes but produced only a modest effect on DOPG-rich membranes. This trend closely parallels the membrane composition-dependent adsorption of FITC-DEX observed in Figure 1. Control experiments confirmed that the initial Laurdan emission spectra were largely comparable among the different membrane compositions before polymer addition (Figure S13), with the exception of membranes containing 30 or 90 mol% DOPG (discussed in the Supporting Information), suggesting that Laurdan experienced a similar local environment across most membrane compositions prior to polymer addition. Moreover, polymer-induced vesicle clustering was similar for all membrane compositions (Figure S14), and the ΔGP values remained essentially unchanged in the presence of 500 mM NaCl (Figure S15), indicating that the observed membrane responses in Figure 3C are neither a consequence of differential crowding nor primarily driven by electrostatic interactions. The strong correlation between polymer adsorption (Figure 1C), multilamellar membrane formation (Figure 2), and GP changes (Figure 3C) suggests that DEX-induced membrane restructuring is closely coupled to changes in the interfacial physicochemical environment sensed by Laurdan.

For comparison, PEG exhibited markedly different behavior (Figure 3D). Across membranes containing 10 mol%+90 mol% DOPC to 90 mol% DOTAP+10 mol% DOPC, PEG induced only minimal increases in GP despite exhibiting composition-dependent membrane adsorption similar to that of DEX (Figure 1D). Given that PEG generated considerable crowding-induced vesicle clustering, these small GP increases are consistent with depletion-induced removal of bulk or loosely bound interfacial water,^44^ which lies largely outside Laurdan’s sensing volume. Interestingly, PEG decreased the GP value of membranes containing 30 or 90 mol% DOPG, indicating increased membrane fluidity - an effect examined further below (Figures 4 and 5). Finally, PEG with molecular weights of 8 and 10 kDa produced nearly identical GP changes to the SUVs (Figure S16), confirming that again, the distinct membrane responses induced by PEG and DEX cannot be attributed simply to differences in polymer molecular weight.

**Figure 4.**
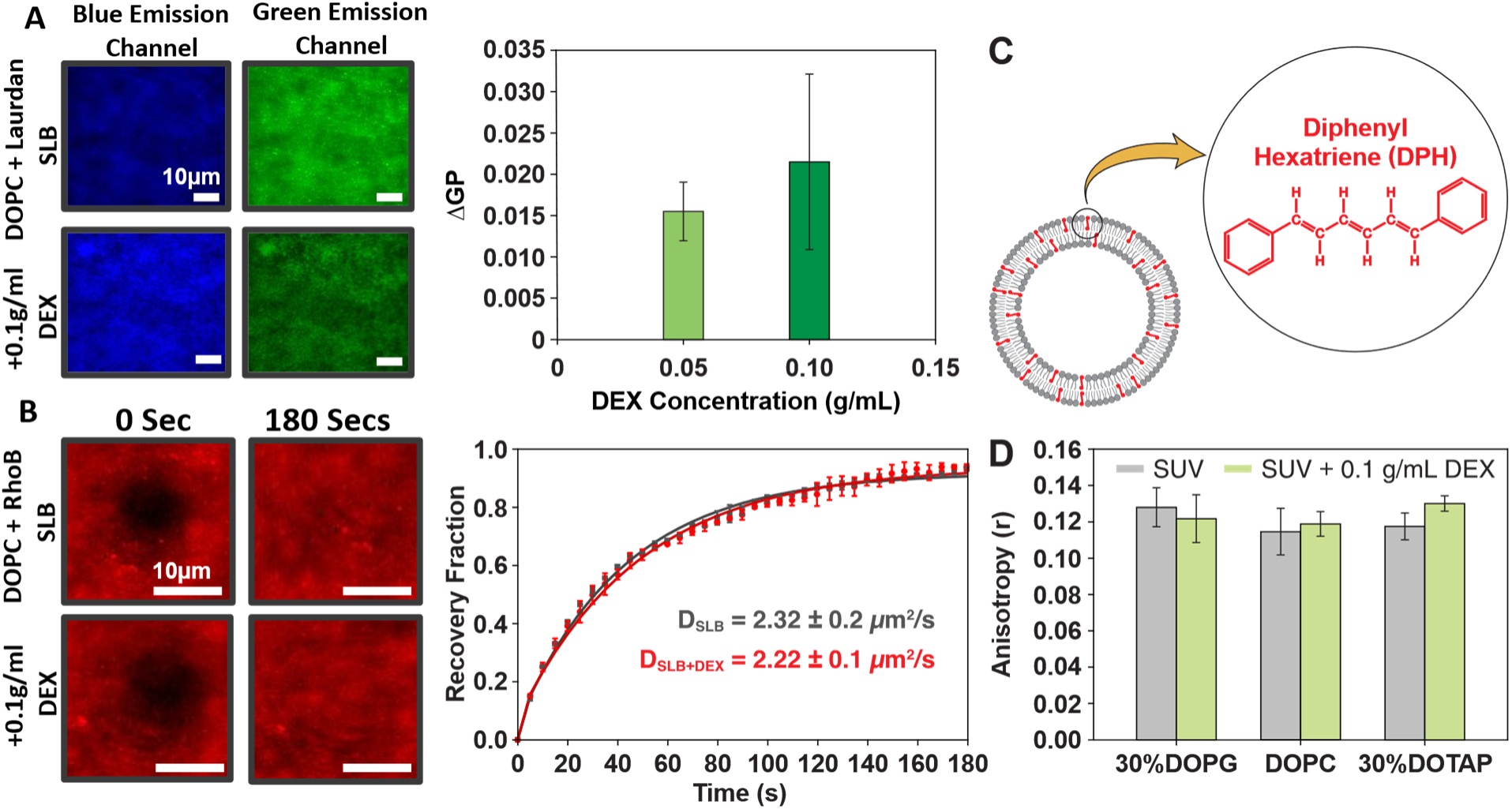
DEX binding does not significantly alter lipid mobility or acyl chain packing. (A, left) TIRF images of SLBs composed of 99.5 mol% DOPC and 0.5 mol% Laurdan before (top) and after (bottom) addition of DEX. Images were collected in the blue emission channel (420–460 nm, left) and green emission channel (470–530 nm, right) (A, right) The bar plot shows ΔGP, calculated from the intensities of the two channels, following addition of 0.05 and 0.1 g/mL DEX. (B, left) FRAP images of SLBs composed of 99.9 mol% DOPC and 0.1 mol% RhoB-PE before photobleaching and 180 s after recovery in the absence (top) and presence (bottom) of 0.1 g/mL DEX. (B, right) Corresponding FRAP recovery curves without DEX (black) and with DEX (red). Symbols represent experimental data, while solid lines represent single-exponential fits. (C) Schematic of DPH incorporated into the hydrophobic region of a lipid bilayer. (D) Fluorescence anisotropy of SUVs containing 0.5 mol% DPH and either 30 mol% DOPG+69.5 mol% DOPC, 99.5 mol% DOPC, or 30 mol% DOTAP+69.5 mol% DOPC in the absence and presence of 0.1 g/mL DEX. Error bars represent standard deviations from four independent experiments performed using four separate SLBs in (A), three independent experiments performed using three separate SLBs in (B), and three independent experiments performed using two separate SUV preparations in (D).

**Figure 5.**
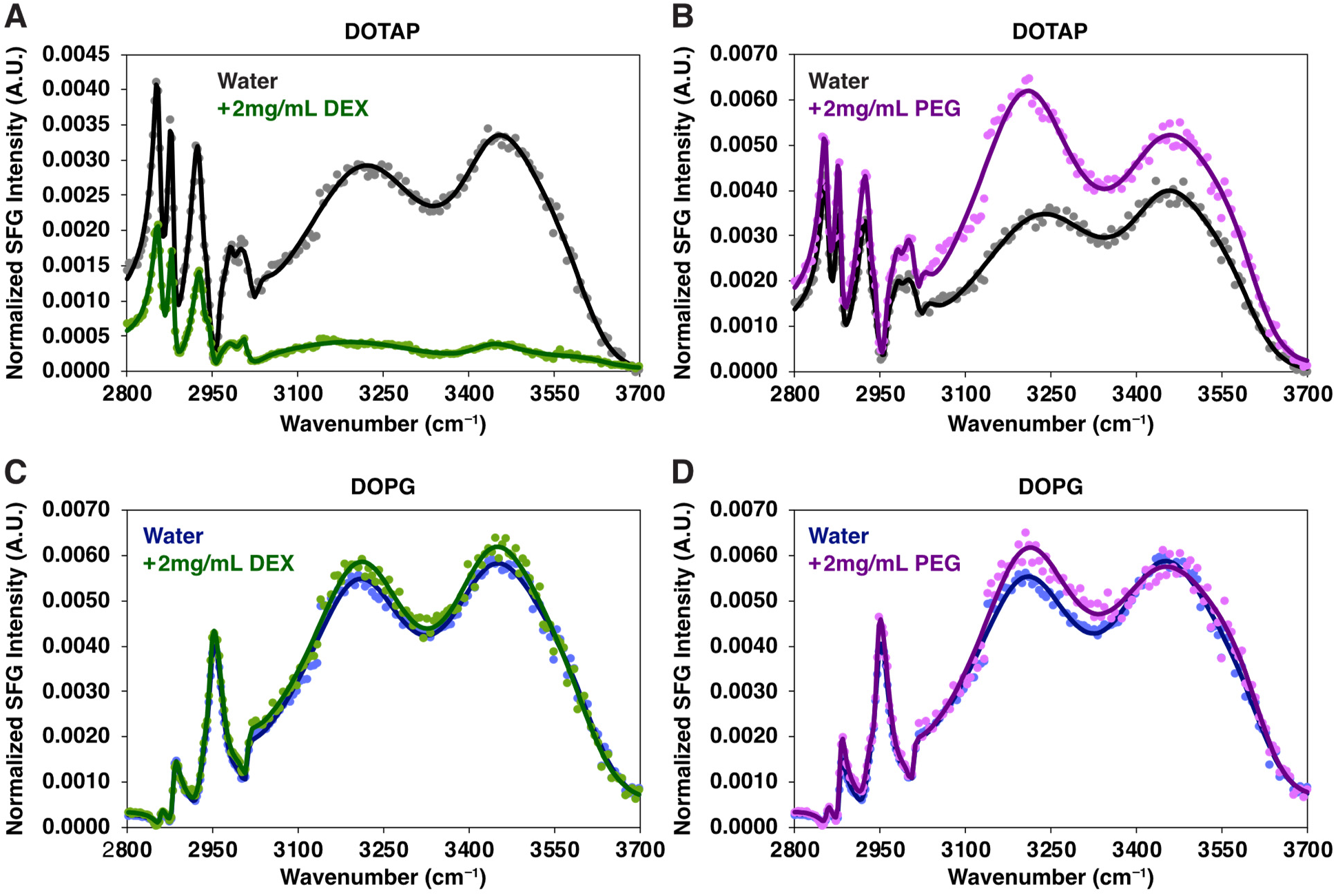
VSFS spectra of DOTAP and DOPG monolayers in the absence and presence of DEX and PEG. (A,B) Spectra of 100 mol% DOTAP monolayers in the absence and presence of 2 mg/mL DEX (A) or PEG (B) in the aqueous subphase. (C,D) Spectra of 100 mol% DOPG monolayers in the absence and presence of 2 mg/mL DEX (C) or PEG (D) in the aqueous subphase. The aqueous subphase is 18 MΩ ultrapure water, and the surface pressure of the lipid monolayer was kept at 17 mN/m for all measurements. The solid circles represent VSFS data points, and the solid lines are fits to the data. All spectra were taken with the ssp polarization combination.

Because both interfacial dehydration and increased lipid acyl-chain packing can contribute to an increase in Laurdan GP (Figure 3C),^38–41^ we next sought to determine which factor primarily accounts for the DEX-induced changes. To probe lipid packing, we first examined whether DEX binding alters lipid lateral mobility (Figure 4A and 4B). SLBs containing 30 mol% DOTAP + 69.5 mol% DOPC and doped with 0.5 mol% Laurdan were prepared, and Laurdan GP values were measured by TIRFM in the absence and presence of DEX (see Methods in the SI for details). Consistent with the bulk measurement (Figure 3C), DEX increased the Laurdan GP of the SLB (Figure 4A), although the magnitude of the increase was smaller than that observed in bulk fluorimetry. This difference is likely owing to the lower lipid content and the absence of membrane curvature in the SLB geometry. To determine whether the increase in GP was accompanied by altered lipid lateral mobility, we performed fluorescence recovery after photobleaching (FRAP) measurements on the SLB in the absence and presence of 0.1 g/mL DEX (Figure 4B). No significant change in the lipid diffusion coefficient was observed following DEX addition. Although FRAP cannot directly quantify acyl-chain packing, lipid packing is generally associated with changes in lateral diffusion.^45, 46^ Therefore, the DEX-induced increase in Laurdan GP is unlikely to originate from substantial changes in membrane dynamics and packing.

To further probe the organization of the hydrophobic core, we employed 1,6-diphenyl-1,3,5-hexatriene (DPH), a well-established fluorescence anisotropy probe that partitions into the acyl-chain region of lipid bilayers and reports on hydrocarbon chain order (also known as membrane microviscosity, Figure 4C).^47, 48^ DPH fluorescence anisotropy measurements of SUVs in the absence and presence of DEX showed only minimal changes (Figure 4D), indicating negligible perturbation of acyl-chain order. This behavior was consistent for DOTAP+DOPC, pure DOPC, and DOPG+DOPC vesicles. Together, the FRAP and DPH anisotropy measurements indicate that DEX does not significantly alter lipid lateral mobility or acyl-chain packing. Combined with the Laurdan GP results (Figures 3C), these findings suggest that the primary effect of DEX is modulation of membrane interfacial hydration rather than changes in lipid packing. PEG induced only a slight increase in DPH anisotropy across the membrane compositions examined (Figure S17), indicating only minimal increase in hydrocarbon chain order. This observation parallels the relatively small positive ΔGP values observed for most membrane compositions (Figure 3D). Notably, for SUVs with high DOPG concentrations, PEG markedly decreased the Laurdan GP (Figure 3D), whereas no corresponding change in DPH anisotropy was detected (Figure S17). These results indicate that the decrease in Laurdan GP is primarily attributable to increased interfacial hydration rather than a disruption in acyl-chain packing (see Figure 5 for additional evidence).

To probe polymer-induced perturbations of lipid headgroups at the molecular level, we employed vibrational sum frequency spectroscopy (VSFS) to characterize pure DOTAP and DOPG monolayers at the air-water interface. The spectra were recorded over frequency ranges corresponding to the acyl-chain portion of the lipid molecules and included the adjacent interfacial water structure. To examine polymer effects, PEG or DEX was introduced into the aqueous subphase. Figure 5A and 5B demonstrate the spectra of DOTAP, in the absence and presence of 2 mg/mL DEX or PEG in the subphase. At this concentration, neither polymer generated detectable SFG signals in the absence of lipids (Figure S18). In the absence of polymer, the water molecules are initially well-aligned by the net positive charge on the DOTAP monolayer (Figure 5A).^49, 50^ Upon addition of DEX, the intensity of the two main water peaks around 3200 cm^-1^ and 3400 cm^-1^ decreased substantially (Table S2, the fitted oscillator strengths retained the same sign). Control experiments ruled out salt contamination in the polymer samples (Table S1), and the suppression of the O– H response persisted in the presence of 150 mM NaCl (Figure S19), demonstrating that the spectral changes are not driven by ionic effects.

The apparent intensity decrease in the C–H stretching region arises primarily from interference with the O– H resonances, which was reported previously with DOTAP lipid monolayers.^49^ After fitting the spectra, the oscillator strengths of the lipid C-H vibrations (2856 cm^-1^ and 2880 cm^-1^) in the acyl chains exhibited only minimal changes (Table S2). Specifically, we observed a slight decrease (12%) in the oscillator strength ratio for the 2880 cm^-1^(CH_3,sym_) and 2856 cm^-1^ (CH_2, sym_) peaks (*I*_CH3,sym_/*I*_CH2,sym_), indicating a slight reduction in acyl-chain ordering.^51^ Thus, despite the pronounced changes in the O–H stretching region, DEX minimally perturbs lipid packing, matching the results from the FRAP and DPH anisotropy measurements (Figure 4). In contrast, DEX increased the oscillator strength of the choline symmetric C-H stretch at 2983 cm^-1^ by more than fivefold, demonstrating a substantial reorganization of the lipid headgroup environment. Because VSFS intensity depends on both molecular orientation and number density, the reduced O–H response likely reflects a combination of decreased interfacial hydration and changes in water ordering. Together with the Laurdan measurements, these results strongly support DEX-induced dehydration of the DOTAP lipid headgroup region as the dominant effect, although contributions from altered orientational ordering of interfacial water cannot be completely excluded. Notably, unlike bulk membrane dehydration,^44^ this local dehydration is not accompanied by a measurable increase in lipid packing. One possible explanation is that molecular moieties of adsorbed DEX chains occupy the inter-headgroup space and thereby prevent tighter lateral packing of the lipids.^52, 53^

PEG produced a markedly different response (Figure 5B). Upon addition of PEG to the subphase, we observed an opposite signal change in the O-H stretch region of the DOTAP lipids: PEG induced a notable increase in the water peaks, and the lowest wavenumber O-H band red shifted from 3229 cm^-1^ (no PEG) to 3214 cm^-1^ (with PEG). The red shift indicates a more strongly hydrogen-bonded interfacial water network.^54–56^ Although some level of crowding-induced dehydration cannot be excluded, its contribution appears considerably smaller than the PEG-induced enhancement of water ordering. This result aligns with the fluorescence spectroscopy results, where a small increase in the Laurdan GP value was only observed with PEG concentration as high as 0.1 g/mL (Figure 3D). In the CH-stretch region, PEG caused only a slight increase (∼10%) in the *I*_CH3,sym_/*I*_CH2,sym_, indicating minimal changes in the acyl-chain ordering. The choline C–H modes also exhibited much smaller intensity changes than observed for DEX, although the asymmetric choline C-H stretch displayed a noticeable red shift of approximately 6 cm^-1^. This shift suggests that PEG perturbs the local chemical environment through directly interacting^57^ with the choline headgroup without inducing any pronounced dehydration as observed for DEX.

Figure 5C and 5D show the corresponding VSFS spectra of DOPG monolayers. At 2 mg/mL, DEX produced only minor spectral changes within the experimental uncertainty (Figure 5C). Increasing the DEX concentration to 10 mg/mL resulted in a measurable decrease in O–H stretching intensity (Figure S20), matching well with the weak membrane adsorption and modest Laurdan GP increases observed for DOPG-rich membranes (Figures 1C and 3C). In the C–H stretching region, DEX increased the *I*_CH3,sym_/*I*_CH2,sym_ by approximately 50%, indicating a modest increase in acyl-chain order. In contrast, PEG induced a small, but consistent, increase in O-H stretching intensity at 2 mg/mL (Figure 5D), with no additional enhancement at 10 mg/mL (Figure S21). Unlike the response observed for DOTAP in Figure 5B, this increase was not accompanied by a frequency shift of the O-H bands. Together with the decrease in Laurdan GP for DOPG-rich membranes (Figure 3D), these results suggest that PEG binding increases hydration within the PG headgroup region.

Collectively, the results presented in Figures 1–5 demonstrate that lipid headgroup chemistry, exemplified here by DOTAP and DOPG, strongly modulates the interactions between lipid membranes and hydrophilic polymers such as PEG and DEX. DOTAP enhances polymer binding, whereas DOPG slightly suppresses it. Although both polymers exhibit similar lipid composition-dependent binding trends, the consequences of binding for membrane structure and interfacial properties differ markedly. Specifically, DEX binding substantially perturbs the hydration shell surrounding the DOTAP headgroup through dehydration, thereby promoting large-scale membrane reorganization and multilamellar membrane formation. In contrast, DEX binding induces only modest dehydration of the DOPG hydration layer. PEG, on the other hand, exerts nearly opposite effects on lipid headgroup hydration. On DOTAP-containing membranes, PEG increases the hydrogen-bonding order of the interfacial water while largely preserving the overall hydration level. On DOPG-containing membranes, PEG increases headgroup hydration with little change in the hydrogen-bonding order of the interfacial water.

To gain insight into the thermodynamic origins of these distinct polymer–membrane interactions, we performed the binding experiments described in Figure 1 over a temperature range of 20–50 °C (Figures 6A and 6B). For each condition, the fluorescent polymers and SLBs were equilibrated for 1 h before fluorescence imaging. Moreover, to minimize potential temperature-dependent photophysical effects on the fluorophores, all TIRFM binding measurements were performed using polymers at near-single-molecule concentrations. As the temperature increased, PEG and DEX exhibited opposite binding behaviors on DOTAP-containing SLBs: DEX binding increased substantially (Figure 6A, dark green), whereas PEG binding decreased (Figure 6B, dark purple). In contrast, on DOPG-containing SLBs, DEX binding also increased prominently with temperature (Figure 6A, light green), whereas PEG binding exhibited little temperature dependence (Figure 6B, light pink). Control experiments further confirmed that polymer– polymer interactions remained essentially unchanged over this temperature range (Figure S22).

**Figure 6.**
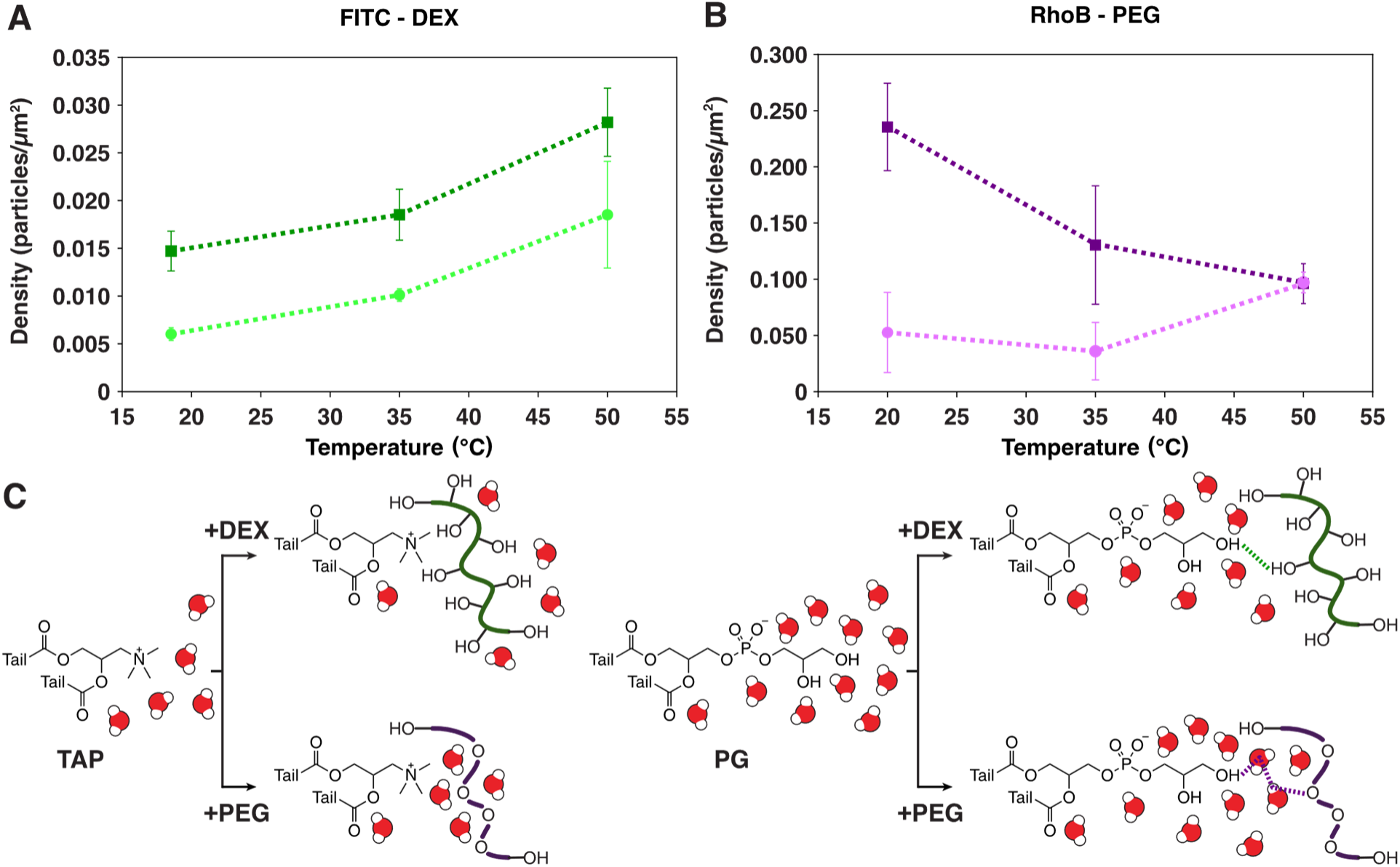
Temperature-dependent binding of DEX and PEG on SLBs containing DOTAP or DOPG. (A) Particle density of surface-bound FITC–DEX on SLBs composed of 30 mol% DOTAP+70 mol% DOPC or 30 mol% DOPG+70 mol% DOPC as a function of temperature. (B) Particle density of surface-bound RhoB–PEG on the same SLBs as a function of temperature. Error bars represent standard deviations from six measurements obtained from two independent SLB preparations. (C) Schematic illustration summarizing the proposed mechanism of how the interfacial hydration layer regulates the interactions between DEX and PEG with DOTAP- and DOPG-containing membranes.

The distinct temperature dependences of PEG and DEX binding suggest fundamentally different thermodynamic origins for their interactions with lipid membranes (Figure 6C). Binding of DEX to both DOPG- and DOTAP-containing SLBs induces dehydration of the close hydration layer surrounding the lipid headgroups (Figures 3 and 5). The release of these hydration water molecules contributes favorably to the entropy of binding. Because DEX binding increases markedly with temperature for both lipid compositions, these observations suggest that DEX–membrane interactions are predominantly entropy driven.^58, 59^ In contrast, PEG binding to DOTAP-containing SLBs induces a red shift of the choline CH_3_ stretching mode while increasing the ordering of interfacial water, without significantly reducing the overall hydration level (Figure 5). Moreover, PEG binding becomes substantially weaker as the temperature increases. Together, these observations indicate that PEG interacts with the TAP headgroup through direct hydrophobic interactions, which become less favorable at elevated temperatures and are therefore associated with a comparatively small favorable entropy contribution.^60^ For PEG binding to DOPG-rich SLBs, the interface remains highly hydrated, with an increased population of hydration water molecules (Figures 3 and 5). In addition, the binding exhibits little temperature dependence. We therefore propose that PEG interacts with the PG headgroup primarily through hydrogen bonding mediated by the interfacial hydration layer. In this case, weakening of hydrogen bonds at higher temperature^61^ may be offset by the accompanying increase in entropy (Figure S23), resulting in a binding free energy that is relatively insensitive to temperature.

Lastly, we seek to understand the molecular mechanism by which lipid headgroup chemistry regulates polymer-membrane interactions. We think that DOTAP promotes polymer binding because its headgroup exhibits stronger characteristics of hydrophobic hydration, resulting in interfacial water molecules that are more weakly associated with the headgroup than those surrounding DOPG.^26,27^ Consequently, these water molecules are more readily replaced or reorganized upon polymer binding, contributing to the favorable entropic driving force for DEX–DOTAP membrane interactions and, we propose, facilitating direct hydrophobic contact during PEG–DOTAP membrane interactions. In contrast, the more strongly bound hydration shell surrounding the DOPG headgroup opposes these processes, leading to weaker polymer– membrane interactions. Having established that electrostatic interactions are not the dominant factor (Figure S4, S15, S19), we next validate this headgroup hydration-based mechanism using phosphatidylethanolamine (PE) and phosphatidic acid (PA) (Figure 7). PE headgroups possess greater hydrogen-bonding capability than PC and therefore maintain a more strongly bound hydration shell,^62^ and correspondingly, DEX-induced Laurdan GP changes decreased with increasing PE content in PC membranes. Likewise, although PA carries a negative charge similar to PG, it contains only a phosphate group and lacks the two additional hydroxyl groups present in PG, resulting in a more weakly bound hydration shell than PG.^63, 64^ Supporting our model, DEX induced larger GP changes in PA-containing than PG-containing membranes. Therefore, these observations suggest that lipid headgroup hydration, rather than lipid charge, is a key determinant of uncharged hydrophilic polymer–membrane interactions and that this mechanism may be broadly applicable to biologically relevant membrane compositions beyond those investigated here.

**Figure 7.**
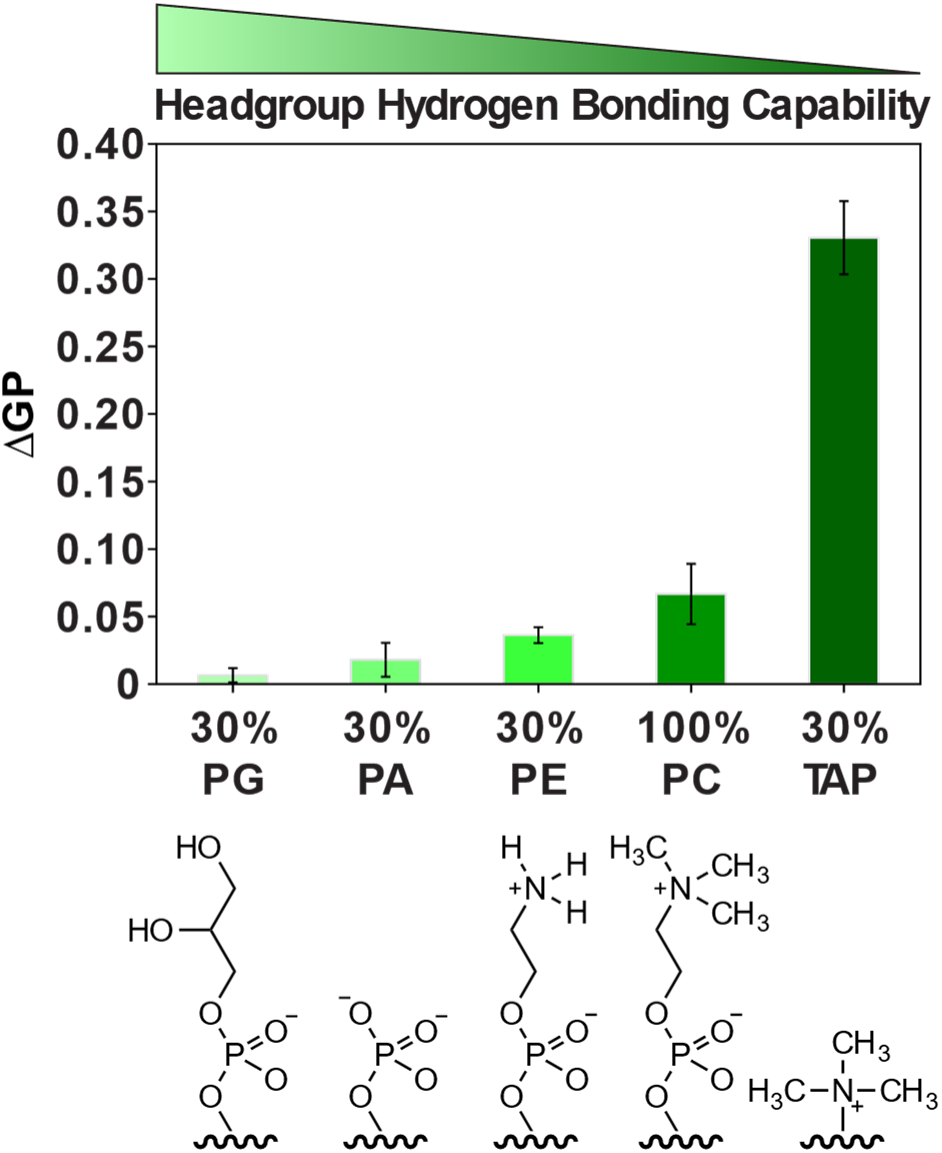
Effect of lipid headgroup chemistry on dextran-induced changes in Laurdan GP. ΔGP values measured following addition of 0.1 g/mL DEX to SUVs containing 0.5 mol% Laurdan and 99.5 mol% POPC, or 30 mol% POPG, POPA, or POPE, with POPC as the remaining lipid component.

## Discussion

Our results reveal that membrane binding affinity alone does not determine the extent to which hydrophilic polymers perturb membrane interfacial environment and structure. Although both PEG and DEX are considered as weakly associated with lipid membranes, their interactions are governed by distinct physicochemical principles and lead to markedly different consequences. PEG exhibits a higher membrane affinity than DEX across DOPG-, DOPC-, and DOTAP-containing membranes (Figure 1E, Figure 6A&6B) and induces stronger crowding mediated vesicle clustering (Figure S10 and S13), which are in line with previous reports.^9, 11^ Surprisingly, despite its stronger membrane association, PEG produces substantially smaller perturbations to the hydration shell surrounding the lipid headgroups and less membrane remodeling than DEX. These contrasting behaviors likely originate from the fundamentally different hydroxyl-group densities and hydration characteristics of the two polymers.

DEX, as a representative neutral polysaccharide, possesses a high density of hydroxyl groups that readily form hydrogen bonds with both water molecules and neighboring hydroxyl groups. Previous studies have shown that dextran exhibits upper critical solubility temperature behavior under appropriate solution conditions,^65^ indicating that polymer-polymer interactions become increasingly favorable as the enthalpic gain from intermolecular hydrogen bonding outweighs the accompanying entropic penalty associated with restructuring hydration water. This behavior suggests that dextran can remain strongly hydrated while maintaining a relatively compact hydration shell. Upon membrane binding, DEX hydroxyl groups can directly form hydrogen bonds with lipid headgroups, partially replacing lipid-associated hydration water. A similar mechanism has been proposed for glucose-lipid interactions.^52^ As the hydration layer surrounding membrane-bound DEX is expected to be relatively thin, as well as structurally disordered,^52, 66^ making only a limited contribution to the Laurdan and VSFS signals.^55^

In contrast, PEG contains a substantially lower density of hydroxyl groups and exhibits some level of hydrophobic characteristics due to its ethylene moieties.^67, 68^ These chemical features render PEG a well-known lower critical solubility temperature polymer,^69, 70^ reflecting the increasing thermodynamic favorability of dehydrating the polymer upon heating as the entropy gained from releasing hydration water overcomes the enthalpic loss from weakened intermolecular interactions. Indeed, PEG is generally considered to possess a relatively extended and well-ordered hydration layer.^71–73^ Previous studies further support this distinction by showing that, although PEG and DEX produce comparable macromolecular crowding effects, PEG possesses a substantially larger hydration shell and correspondingly lowers the water potential to a greater extent.^66^ Accordingly, PEG likely associates with lipid membranes primarily through hydrogen-bond acceptance by its ether oxygens together with interactions involving its amphiphilic backbone, while largely preserving the surrounding hydration layer. At DOTAP-containing membranes, PEG binding enhances the ordering of interfacial water without substantially reducing the overall hydration level, possibly through the formation of a shared hydration network between PEG and the lipid headgroups. In contrast, at PG-containing membranes, the bulky hydration shell surrounding PEG may increase the thickness of the interfacial hydration layer separating the polymer from the lipid headgroups. Collectively, these results suggest that DEX primarily remodels the membrane interface by replacing lipid hydration water, whereas PEG largely preserves and reorganizes the existing hydration layer. More broadly, these distinct hydration characteristics may also contribute to the thermodynamic driving forces governing PEG– DEX aqueous two-phase separation and the differential partitioning of biomolecules between the two phases, further supporting the emerging view that water plays an active and thermodynamically important role in biomacromolecular phase separation.^74, 75^

## Acknowledgments

This work was supported by the Stony Brook University Startup Research Fund (to S.S.) and NIH NIGMS 1R35GM162491-01(to S.S.). S.B. was supported by the Stony Brook Chemistry– Biology Training Program (CBTP), funded by the National Institutes of Health Training Grant T32GM136572. We thank E. London and D. P. Raleigh for sharing the fluorescence spectrometer. We thank P. S. Cremer for sharing the VSFS instrumentation. We acknowledge Dr. Fang Lu at Brookhaven National Laboratory (BNL) for assistance with zeta potential measurements. We thank M. R. Fernandez for valuable discussions.

## Author contributions

S.S. and S.B. conceived the study and designed the experiments. S.B. performed the experiments and analyzed the experimental data, except for the VSFS experiments. S.S. performed the VSFS measurements and analyzed the corresponding data. D.N. assisted with AFM measurements and data analysis. J.L. contributed to data interpretation and prepared figures for the manuscript. S.B. and S.S. wrote the manuscript. All authors discussed the results and approved the final manuscript.

## Data availability

All data supporting the findings of this study are available within the paper and its Supporting Information. Additional data are available from the corresponding author upon reasonable request.

