## Supplemental Materials for "Lipid Headgroup Hydration Regulates Distinct Remodeling of the Membrane Interface by Polyethylene Glycol and Dextran"

### 1.Materials

Lipids including 1-palmitoyl-2-oleoyl-sn-glycero-3-phosphocholine (POPC), 1,2-dioleoyl-sn-glycero-3-phosphocholine (DOPC), 1-palmitoyl-2-oleoyl-sn-glycero-3-phospho-(1'-rac-glycerol) (POPG), 1,2-dioleoyl-sn-glycero-3-phospho-(1'-rac-glycerol) (DOPG), 1,2-dioleoyl-3-trimethylammonium-propane (DOTAP), 1-palmitoyl-2-oleoyl-sn-glycero-3-phosphoethanolamine (POPE), 1-palmitoyl-2-oleoyl-sn-glycero-3-phosphate (POPA), and 1,2-dioleoyl-sn-glycero-3-phosphoethanolamine-N-(lissamine rhodamine B sulfonyl) (ammonium salt) (RhoB-PE) were purchased from Avanti Polar Lipids (Alabaster, AL) and supplied in chloroform solution. Laurdan (6-dodecanoyl-2-dimethylaminonaphthalene) and β-BODIPY™ FL C12-HPC(2-(4,4-difluoro-5,7-dimethyl-4-bora-3a,4a-diaza-s-indacene-3-dodecanoyl)-1-hexadecanoyl-sn-glycero-3-phosphocholine) were purchased from Thermo Fisher Scientific (Invitrogen) as powders and dissolved in chloroform to prepare stock solutions. Throughout this work, β-BODIPY™ FL C12-HPC is referred to as BODIPY-PC. 1,6-diphenyl-1,3,5-hexatriene (DPH) was purchased from Sigma-Aldrich in powder form and dissolved in chloroform to prepare stock solutions.

Poly (ethylene glycol) with average molecular weights of 8 and 10 kDa (PEG 8K and PEG 10K; BioUltra, molecular biology grade) was purchased from Sigma-Aldrich. Dextran with an average molecular weight of 9–11 kDa (DEX 10K), derived from *Leuconostoc mesenteroides* and containing approximately 95% linear and 5% branched structures, was also purchased from Sigma-Aldrich. Throughout this work, PEG 8K and DEX 10K are referred to as PEG and DEX, respectively, unless otherwise noted.

Fluorescein isothiocyanate-labeled dextran (FITC-DEX; average molecular weight 10 kDa), derived from *Leuconostoc mesenteroides* strain B512, was purchased from MilliporeSigma. According to the manufacturer, FITC is randomly conjugated to the hydroxyl groups of dextran through a stable thiocarbamoyl linkage at a frequency of 0.003–0.02 mol FITC per mol glucose, resulting in minimal modification of the native dextran backbone. Rhodamine B-labeled poly(ethylene glycol) (RhoB-PEG average molecular weight 8 kDa; purity ≥90%) was purchased from Biopharma PEG. RhoB-PEG contains a single Rhodamine B fluorophore conjugated to one terminus of each PEG chain. Throughout this work, fluorescein isothiocyanate-labeled dextran and rhodamine B-labeled poly (ethylene glycol) are referred to as FITC-DEX and RhoB-PEG, respectively. All polymers were dissolved in ultrapure water (18.2 MΩ·cm, Milli-Q) prior to use.

Sterile-filtered 1× phosphate-buffered saline (PBS, pH 7.4) was obtained from Thermo Fisher Scientific and used for supported lipid bilayer preparation and imaging experiments. Chloroform (ACS grade, ≥99.8%, Thermo Fisher Scientific) was used for lipid and fluorophore stock preparation. Ultrapure water (18.2 MΩ·cm at 25 °C) produced using a Milli-Q IQ 7000 water purification system (MilliporeSigma) was used for all solution preparation.

### 2. Methods

#### Glass Cleaning Procedure

Glass coverslips were first sonicated in a 1 wt% sodium dodecyl sulfate (SDS) solution for 1 hour. The coverslips were then transferred to 70% ethanol and sonicated for an additional 1 hour. After cleaning, the coverslips were rinsed thoroughly with ultrapure water (18.2 MΩ·cm at 25 °C) and dried under a stream of nitrogen gas. Finally, the coverslips were plasma cleaned using a Harrick Plasma Cleaner (Ithaca, NY, USA). Plasma treatment was carried out under vacuum until the chamber pressure dropped below ~1 torr and a characteristic pink glow discharge was observed. The glass substrates were exposed to plasma for 5 minutes before depositing SUVs.

#### Preparation of Small Unilamellar Vesicles (SUV)

SUVs were prepared using standard freeze-thaw and extrusion techniques[1, 2]. Lipids were mixed in the desired ratio in chloroform. The chloroform was subsequently removed by drying under a stream of nitrogen gas followed by placing the lipids under vacuum in a desiccator for at least 2 hours. The dried lipids were rehydrated in 1× PBS at pH 7.4 to a final lipid concentration of 1 mg/mL and vortexed briefly. The rehydrated lipids were then subjected to ten freeze-thaw cycles using liquid nitrogen and a 45 °C water bath. After freeze-thawing, the lipids were extruded twenty-one times through a track-etched polycarbonate membrane with 100 nm pores (Whatman, Florham Park, NJ). Dynamic light scattering (DLS) confirmed the monodispersity and stability of the vesicles.

For fluorometry measurements, SUVs containing Laurdan or DPH were prepared using the same procedure. In these cases, the dried lipids were rehydrated in ultrapure water (18.2 MΩ·cm at 25 °C) instead of PBS to a final lipid concentration of 0.5 mg/mL. All other preparation steps remained unchanged.

#### Formation of Supported Lipid Bilayer (SLB)

SLBs were prepared by the vesicle fusion method[3]. A six-channel ibidi Sticky-Slide VI 0.4 (ibidi GmbH, Germany) was attached to the plasma-cleaned glass coverslips described above to create six independent imaging chambers. SUVs were first prepared at a lipid concentration of 1 mg/mL in 1× PBS buffer. The SUV solution was then diluted two-fold with ultrapure water (18.2 MΩ·cm at 25 °C) to a final lipid concentration of 0.5 mg/mL. A 100 µL aliquot of the diluted SUV solution was introduced into the channel and allowed to incubate for 30 minutes. After incubation, excess vesicles were removed by washing with copious 1× PBS buffer.

Dilution of the SUV suspension prior to bilayer formation was necessary to minimize the presence of unfused vesicles on the surface. Any remaining unfused vesicles were removed during the washing step following incubation (as demonstrated by AFM measurements in Figure S9).

All microscope imaging measurements described herein were conducted at room temperature (22 ± 0.5 °C), except for the temperature-dependent experiments.

#### Preparation of Polymer Solutions

Polymer solutions were prepared using ultrapure water (18.2 MΩ·cm at 25 °C) nd used within 3 days of preparation. Polymer powder was weighed into a glass vial and dissolved in water by vortexing. If necessary, the solution was sonicated until the polymer was completely dissolved. Prior to use, all polymer solutions were filtered through a 0.22 μm syringe filter. The excluded volume of the polymer was taken into consideration when calculating the amount of polymer and water required to obtain the desired final concentration. To prepare a 0.1 g/mL polymer solution, approximately 0.12 g of polymer was weighed into a vial and initially dissolved in 800 μL of ultrapure water. After complete dissolution by vortexing and sonication, the solution volume was measured, and additional ultrapure water was added as needed to obtain a final polymer concentration of 0.1 g/mL.

Fluorescently labeled polymer solutions were prepared differently. The entire polymer sample was dissolved in water at the stock concentration provided by the manufacturer. The stock solution was aliquoted into 100 μL portions and stored at −20 °C. Prior to an experiment, a single aliquot was thawed and diluted to the desired concentration using ultrapure water (18.2 MΩ·cm at 25 °C) . The diluted solution was filtered through a 0.22 μm syringe filter before use. Each aliquot was thawed only once, and any remaining solution was discarded after use.

**Microscopy**

TIRF experiments were performed on a motorized inverted microscope (Nikon Eclipse Ti2-E, Nikon Instruments Inc., Melville, NY) ) equipped with a motorized Epi/TIRF illuminator, motorizedSOLA light engine,, Perfect Focus system, and a motorized stage (MS-2000, Applied Scientific Instrumentation, Eugene, OR). A laser enginewith 405, 488, 561 and 640-nm (LUN-F XL) diode lasers were controlled and aligned into a fiber launch. The optical path was then aligned to a 100× 1.49–numerical aperture oil immersion TIRF objective (Nikon). A dichroic beamsplitter (ZT405/488/561/640rpc, Chroma Technology Corp., Bellows Falls, VT) reflected the laser light through the objective lens, and fluorescence images were recorded using an ORCA Fusion BT scientific CMOS camera (C15440, Hamamatsu Photonics, Japan) after passing through a laser-blocking filter (ZET405/488/561/640m-TRF, Chroma Technology Corp., Bellows Falls, VT) and a barrier filter for cleaning up. Laser powers measured at the sample stage were approximately 19 mW (405 nm), 48 mW (488 nm), 51 mW (561 nm), and 68 mW (640 nm), when set at 100% power output. All acquisitions were obtained using NIS-Elements software.

**Fluorescence Imaging and Single Particle Analysis**

Membrane composition-dependent polymer adsorption was investigated using both ensemble fluorescence imaging and single particle imaging.

For ensemble imaging measurements, FITC-DEX and RhoB-PEG were added to supported lipid bilayers at a final concentration of 0.1 μM. FITC-DEX was imaged using the 488 nm laser and RhoB-PEG was imaged using the 561 nm laser. Images were acquired using the microscope configuration described above with 15% laser power. The TIRF images were flattened in Fiji ImageJ prior to analysis using a Gaussian blur filter with a sigma radius of 50 pixels. For each sample, five randomly selected regions were imaged. The average fluorescence intensity of each image was measured using Fiji ImageJ, and the values from the five random regions were averaged to obtain the fluorescence intensity reported for that sample.

For single particle imaging measurements, FITC-DEX was used at a concentration of 10 nM and RhoB-PEG was used at a concentration of 0.6 nM. Representative particle intensity distributions are shown in Figure S1 to demonstrate that the detected fluorescence spots correspond predominantly to a single particle population. FITC-DEX was imaged using the 488 nm laser and RhoB-PEG was imaged using the 561 nm laser. Images were acquired using the same microscope configuration with the laser power increased to

50%. The TIRF images were flattened and background subtractedprior to analysis. Image flattening was performed in Fiji ImageJ using a Gaussian blur filter with a sigma radius of 50 pixels. Background substraction was performed using a rolling ball algorithm with a radius of 50 pixels. Individual particles were identified and counted using the ComDet plugin (version 0.5.5). The approximate particle size was set to 3 pixels, and the intensity threshold was set to 3 standard deviations above the local background. Larger particles were included and segmented during analysis. Oval regions of interest were used for particle identification. The average number of particles per image area was calculated and reported. Each image corresponded to an area of $4.43 \times 10^3$ $\mu m^2$.

**Fluorescence Imaging of Membrane Patches**

Membrane patches were visualized using supported lipid bilayers containing either 0.1 mol% RhoB-PE or 0.5 mol% BODIPY-PC. Unlabeled PEG or DEX was added to the supported lipid bilayers at a final concentration of 0.1 g/mL - 0.3 g/mL. RhoB-PE was imaged using the 561 nm laser and BODIPY-PC was imaged using the 488 nm laser. Images were acquired using the same microscope configuration described above with the laser power set to 5%.

The TIRF images were flattened prior to analysis in Fiji ImageJ using a Gaussian blur filter with a sigma radius of 50 pixels. Line profile analysis was carried out by drawing a line across the membrane patches and measuring the fluorescence intensity as a function of distance. The fluorescence intensity of the patches and the surrounding bilayer was measured from the resulting line profiles.

The average fluorescence intensity of the bilayer was measured before polymer addition. After polymer addition, membrane patches were identified by thresholding the fluorescence images in Fiji ImageJ. The resulting selection was used to measure the average fluorescence intensity of the patches. The selection was then inverted and used to measure the average fluorescence intensity of the surrounding bilayer. The ratio of the patch intensity to the surrounding bilayer intensity was subsequently calculated.

**Laurdan Imaging and Generalized Polarization Analysis**

SLBs containing 0.5 mol% Laurdan were imaged using the microscope configuration described above. Laurdan was excited using a 405 nm laser. The emission was collected in two separate spectral ranges. The blue emission channel (420–460 nm) was used to measure the fluorescence intensity around 440 nm, while the green emission channel (470–530 nm) was used to measure the fluorescence intensity around 490 nm.

Images were acquired before and after polymer addition. Prior to analysis, the images were flattened in Fiji ImageJ using a Gaussian blur filter with a sigma radius of 50 pixels. The average fluorescence intensity of the blue and green channels was measured and used to representthe fluorescence intensities at 440 nm and 490 nm, respectively for calculating the generalized polarization (GP) value [4]:

$$GP = (I_{440} - I_{490}) / (I_{440} + I_{490}) \qquad (1)$$

The change in generalized polarization (ΔGP) was calculated by subtracting the GP value before polymer addition from the GP value after polymer addition and plotted as a function of polymer concentration.

Discussion paragraph on what Laurdan is sensing in this work and the underlying mechanism based on our understanding.

**Fluorescence Recovery After Photobleaching (FRAP)**

The lateral diffusion of lipids in supported lipid bilayers (SLBs) was quantified by fluorescence recovery after photobleaching (FRAP). A circular region of the bilayer was photobleached for 3 min using the 561 nm laser. The average radius of the bleached region was 8.6 μm. Following photobleaching, fluorescence images were acquired every 5 s for 3 min to monitor fluorescence recovery as unbleached fluorophores diffused into the bleached region.

The fluorescence intensity within the bleached region was normalized to three unbleached reference regions on the same SLB, and the recovery fraction, F(t), was calculated as[5, 6]:

$$\mathrm{F(t)} = \frac{F_t - F_0}{1 - F_0} \qquad (2)$$

where $F_t$ is the normalized fluorescence intensity at time t and $F_0$ is the normalized fluorescence intensity immediately after bleaching.

The recovery curves were fitted in MATLAB to a single-exponential function,

$$\mathrm{F(t)} = a\,(1 - e^{-bt}) \qquad (3)$$

where a is the mobile fraction and b is the recovery rate constant. The half-time of recovery was calculated as

$$t_{1/2} = \frac{\ln(2)}{b} \qquad (4)$$

The diffusion coefficient (D) was calculated according to:

$$\mathrm{D} = \frac{\omega^2}{4t_{1/2}} \qquad (5)$$

where the radius of the bleached region is ω. The coefficient of determination ($R^2$) and the root mean square error (RMSE) were used to evaluate the quality of the fits. Diffusion coefficients were obtained from the fitted recovery curves.

**Laurdan Fluorescence Spectroscopy**

SUVs containing 0.5 mol% Laurdan were prepared as described above. Laurdan fluorescence measurements were performed using a Fluorolog-QM spectrofluorometer (HORIBA). Prior to data collection, Rhodamine B was used as a fluorescence standard to verify wavelength calibration. Laurdan was excited at 360 nm and emission spectra were collected from 400 to 550 nm. Spectra were collected before and after polymer addition.

**Spectral Fitting and Generalized Polarization Analysis**

Laurdan emission spectra were normalized to the maximum fluorescence intensity prior to analysis. The normalized spectra were fit using a two-Gaussian model corresponding to emission maxima near 440 nm and 490 nm[7]. The Gaussian function used for fitting was:

$$y = y_0 + A/\,(w \cdot \sqrt{\pi}/\,(4 \ln 2)) \cdot \exp\,(-4 \ln 2 \cdot (x - x_c\, w)^{\,2}) \qquad (6)$$

where ($y_0$) is the baseline offset, (A) is the peak area, ($x_c$) is the peak center, and (w) is the full width at half maximum.

Peak heights were calculated from the fitted parameters according to:

$$Height\ (H) = A/\ (w \cdot \sqrt{\pi}/\ (4 \ln 2)) \qquad (7)$$

The peak heights corresponding to the fitted emission maxima near 440 nm and 490 nm were used to calculate the generalized polarization (GP) according to:

$$GP = (I_{440} - I_{490})/\ (I_{440} + I_{490}) \qquad (1)$$

where ($I_{440}$) and ($I_{490}$) are the fluorescence intensities obtained from the fitted peak heights. GP values were calculated before and after polymer addition. The change in generalized polarization (ΔGP) was calculated by subtracting the GP value before polymer addition from the GP value after polymer addition. The quality of the Gaussian fits was evaluated using the coefficient of determination ($R^2$), adjusted ($R^2$), and reduced chi-square.

**DPH Fluorescence Anisotropy Measurements**

DPH fluorescence anisotropy measurements were performed using the same Fluorolog-QM spectrofluorometer (HORIBA) described above and equipped with automatic excitation and emission polarizers. Samples were excited at 360 nm, and the fluorescence emission spectrum was collected from 380 to 600 nm.

The anisotropy analysis were conducted following previous published procedure[8]. The fluorescence intensities corresponding to the HH, HV, VV, and VH polarization configurations were collected for each sample, where the first letter denotes the orientation of the excitation polarizer and the second letter denotes the orientation of the emission polarizer. The fluorescence intensities were averaged over the wavelength range of 420–450 nm prior to analysis.

The instrumental correction factor (G) was calculated according to:

$$G = \frac{I_{HV}}{I_{HH}} \qquad (8)$$

where $I_{HV}$ and $I_{HH}$ are the average fluorescence intensities measured between 420 and 450 nm.

The steady-state fluorescence anisotropy ((r)) was then calculated according to:

$$r = \frac{I_{VV} - GI_{VH}}{I_{VV} + 2GI_{VH}} \qquad (9)$$

where $I_{VV}$ and $I_{VH}$ are the average fluorescence intensities measured between 420 and 450 nm. The anisotropy values reported were calculated from the average fluorescence intensities within this wavelength range.

**Atomic Force Microscopy (AFM)**

AFM measurements were performed using a Cypher ES Environmental AFM (Oxford Instruments Asylum Research) at Brookhaven National Laboratory. Supported lipid bilayers were prepared as described above on 15 mm round glass coverslips. The coverslips were mounted onto 15 mm AFM specimen metal discs prior to imaging. AFM images were acquired in liquid using tapping mode with photothermal excitation and HQ/Cr-Au BS probes (MikroMasch; nominal spring constant 75 N $m^{-1}$, nominal tip radius <2.8 nm). Bilayers were imaged before and after polymer addition. AFM scans were collected over a 5 × 5 $\mu m^2$ area at a scan rate of 0.5 Hz.

Image analysis was performed using Gwyddion (version 2.70). Height profiles were obtained by drawing line sections across the features of interest and measuring the height as a function of distance.

**Dynamic Light Scattering (DLS)**

DLS measurements were performed using a NanoBrook 90Plus particle size analyzer (Brookhaven Instruments, Holtsville, NY, USA) to characterize SUVs before and after polymer addition. SUV samples were prepared as described above. For each measurement, 67 µL of sample was diluted with 433 µL of the corresponding buffer to a final volume of 500 µL and transferred to a disposable cuvette.

Measurements used to characterize vesicle size before and after polymer addition were performed at 25 °C. Temperature-dependent measurements were performed at 5, 35, and 65 °C.

Each sample was measured in five independent runs. The effective diameter (Z-average) and intensity-weighted particle size distributions were obtained from the instrument software. The average effective diameter was calculated from the five measurements and reported as mean ± standard deviation. Intensity-weighted size distributions were used to compare vesicle size distributions under different experimental conditions.

**Vibrational Sum-Frequency Spectroscopy (VSFS)**

VSFS is a second-order nonlinear optical technique.[9] To perform an experiment, a fixed wavelength visible laser beam (532 nm) and a tunable infrared (IR) beam are overlapped in space and time at the lipid/water interface, and the signal which is generated has a frequency that is the sum of the IR and the visible components. The intensity of the emitted VSFS field is strongly enhanced when the IR frequency is on resonance with a vibrational transition from molecules at the surface.[10] To generate a vibrational spectrum at the air/lipid monolayer/water interface, the intensity of the sum frequency beam is monitored as a function of the IR beam frequency. The intensity of the signal for a particular molecular vibration is proportional to the square of the second-order nonlinear susceptibility of the sample. In the dipole approximation, the second order non-linear susceptibility is non-zero in media that lacked inversion symmetry. As such, VSFS is inherently surface specific.[10]

The VSFS experimental setup (EKSPLA, Lithuania) consisted of a 1064 nm Nd: YAG laser (pulse duration: 30 ps; pulse energy: 40 mJ; maximum repetition rate 50 Hz), which was directed to a harmonic unit (H500). The second harmonic (532 nm) and fundamental beams from the harmonic unit were used to pump an optical parametric generator/difference frequency generator (PG501/DFG) unit. The infrared frequency could be tuned between 900 $cm^{-1}$ and 4000 $cm^{-1}$ and the spectral resolution was < 6 $cm^{-1}$. VSFG experiments at the air/water interface, a PTFE Langmuir trough (NIMA technology, England) equipped with a pressure sensor and controllable barriers was used. The trough had an area of 65 $cm^2$ and the subphase volume was

35 ml. A Langmuir monolayer of lipid molecules was formed at the air/water interface by spreading droplets of lipid solutions in chloroform at a concentration of 1 mg/ml. The volume of each drop was 1.5– 2 µl. VSFS spectra were collected in the absence and presence of 2mg/ml polymer solution in the aqueous subphase. The incident angles of the IR and visible beams were 55 and 60°, respectively, with respect to the surface normal at the air/water interface. The energy of the visible beam at the sample stage was 400 µJ and the infrared beam was 150 µJ at 2880 $cm^{-1}$, and 15 µJ at 1100 $cm^{-1}$. The VSFS signal was collected at 2 $cm^{-1}$ intervals and each data point in a spectrum consisted of an average of 200 laser S13 shots. Each spectrum was taken within 40 minutes or less. The nitrogen purging system protected the four double bonds from oxidation, which was confirmed by the lack of spectral change at 3019 $cm^{-1}$ over the course of a given experiment. Each spectrum presented herein has been taken at least three times.

VSFG spectra from the air/water interface were obtained on lipid monolayers composed of 100 mol% DOTAP or 100 mol% DOPG. The monolayers were studied at a surface pressure of 17 mN/m and 22 °C. To investigate polymer–membrane interactions, PEG or DEX was dissolved directly in the aqueous subphase at the indicated concentrations. For salt experiments, the aqueous subphase contained 150 mM NaCl.

The intensity of the sum frequency signal was calculated according to:

$$I_{SFG} \propto |\chi_{eff}^{(2)}|^2 I_{vis} I_{IR} \qquad (10)$$

where $I_{VIS}$ and $I_{IR}$ are the intensities of the incoming visible and infrared laser beams, respectively. $\chi_{eff}^{(2)}$ represents the second order nonlinear susceptibility, which can be further expressed as:

$$\chi_{eff}^{(2)} = \chi^{(2)}_{NR} + \chi^{(2)}_{R} = \chi^{(2)}_{NR} + \sum_n \frac{A_n}{\omega_{IR} - \omega_n + i\Gamma_n} \qquad (11)$$

where $\chi^{(2)}_{NR}$ and $\chi^{(2)}_{R}$ are the frequency independent non-resonant susceptibility term and the frequency dependent resonant susceptibility term, respectively. $\chi^{(2)}_{R}$ of the $n^{th}$ resonant mode is a function of the oscillator strength, $A_n$, resonant frequency, $\omega_n$, peak width, $\Gamma_n$, and the frequency of the input infrared laser beam, $\omega_{IR}$.[10] All data were taken with the ssp polarization combination, which refers to s – sum frequency, s – visible, and p – infrared electric field polarizations. The spectra have each been normalized to the intensities of the incoming visible and IR beams. The normalized spectra were fit to Eqn. 11 using MATLAB. The fitted oscillator strengths, $A_n$, and the peak widths, Γn, are listed in Table S2 – S6, corresponding to each frequency region of interest. The error bars are standard deviations and were calculated by fitting each spectrum individually.

### Supporting Figures

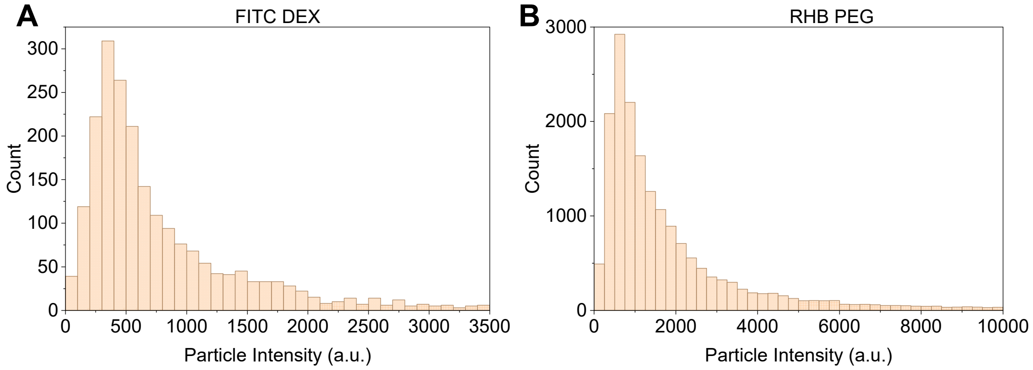

**Figure S1:** Particle intensity distributions of FITC-DEX (10 nM) (left) and RhoB-PEG (0.6 nM) (right) obtained from single-particle imaging experiments on supported lipid bilayers containing 30 mol% DOTAP and 70 mol% DOPC. A total of 2155 FITC-DEX particles (median intensity = 559 a.u.) and 18409 RhoB-PEG particles (median intensity = 1228 a.u.) were analyzed. Both distributions are dominated by low-intensity fluorescent particles with a small tail toward higher intensities, indicating that most detected spots correspond to isolated fluorescent particles, with only a small fraction of brighter spots likely arising from multiple fluorophores, closely spaced particles, or occasional aggregates.

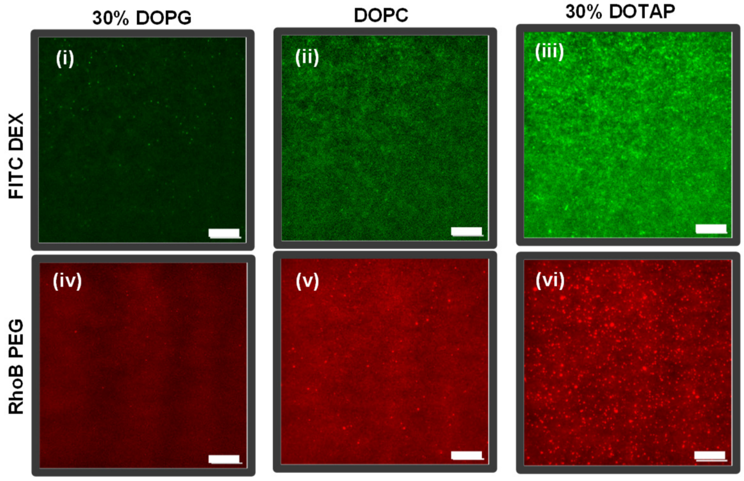

**Figure S2:** Representative TIRF images showing surface adsorption of fluorescent polymers onto SLBs of different compositions. Top row: 0.1 µM FITC-DEX on 70 mol% DOPC + 30 mol% DOPG (i), 100 mol% DOPC (ii), and 70 mol% DOPC + 30 mol% DOTAP (iii). Bottom row: 0.1 µM RhoB-PEG on 70 mol% DOPC + 30 mol% DOPG (iv), 100 mol% DOPC (v), and 70 mol% DOPC + 30 mol% DOTAP (vi). Brighter fluorescence indicates greater polymer adsorption to the membrane. Scale bars, 10 µm.

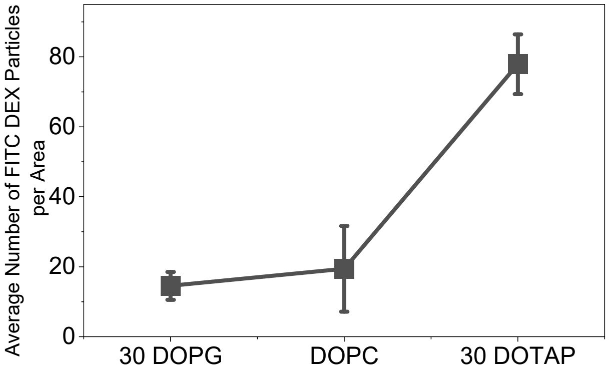

**Figure S3:** Average number of surface-bound FITC-DEX molecules per $4.43 \times 10^3$ μm² imaging area on supported lipid bilayers composed of 70 mol% DOPC + 30 mol% DOPG, 100 mol% DOPC, and 70 mol% DOPC + 30 mol% DOTAP. Measurements were performed using 10 nM FITC-DEX to minimize polymer–polymer interactions. The same trend observed at higher polymer concentrations was still present, with the largest number of bound molecules found on DOTAP-containing membranes. Error bars represent standard deviations from four independent experiments using two separate sets of SUVs.

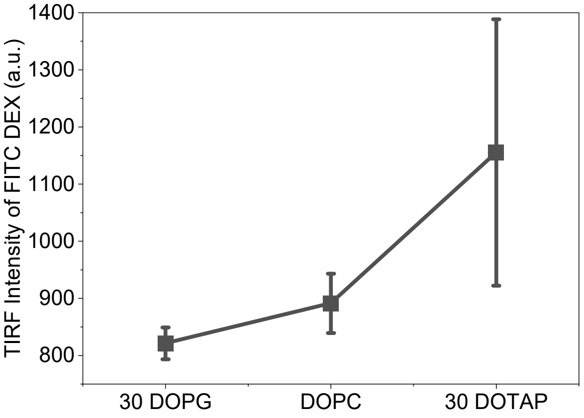

**Figure S4**: TIRF intensity of surface-bound FITC-DEX on supported lipid bilayers composed of 70 mol% DOPC + 30 mol% DOPG, 100 mol% DOPC, and 70 mol% DOPC + 30 mol% DOTAP in the presence of 150 mM NaCl. Measurements were performed using 0.1 μM FITC–DEX to test the role of electrostatic interactions in polymer adsorption. A similar trend was still observed under salt conditions, suggesting that the enhanced adsorption of DOTAP-containing membranes is not mainly driven by simple electrostatic attraction. Error bars represent standard deviations from six independent experiments using three separate sets of SUVs.

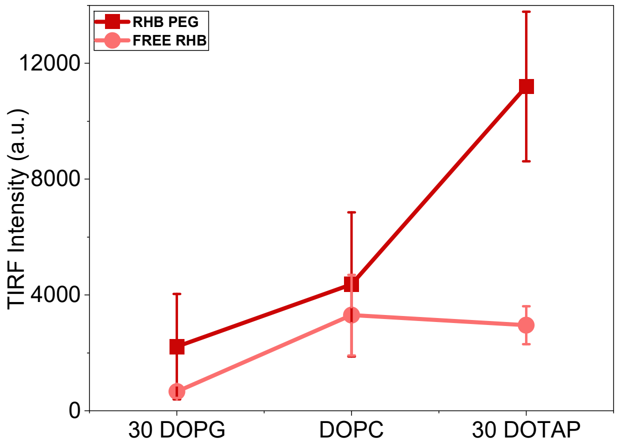

**Figure S5:** Membrane adsorption of free Rhodamine B (RhoB) on supported lipid bilayers composed of 100 mol% DOPC, 70 mol% DOPC + 30 mol% DOPG, and 70 mol% DOPC + 30 mol% DOTAP. Unlike RhoB-PEG, free RhoB exhibited minimal membrane adsorption and showed little dependence on membrane composition. Error bars represent standard deviations from two independent experiments using two separate SUV preparations.

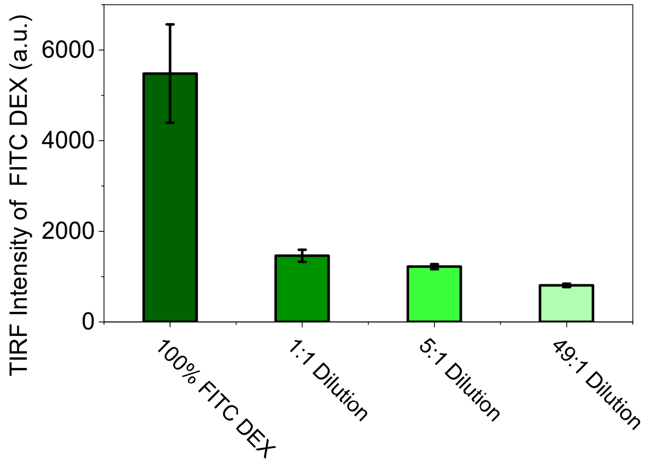

**Figure S6:** Competition binding assay between FITC-DEX and PEG on supported lipid bilayers composed of 70 mol% DOPC + 30 mol% DOTAP. The total polymer concentration was maintained at 0.5 µM, while the fraction of FITC-DEX (0.5 µM at the 100% FITC-DEX condition) was progressively decreased and replaced with unlabeled PEG. Surface-bound FITC-DEX was quantified by TIRF microscopy. No aqueous two-phase separation was observed at this polymer concentration. FITC fluorescence decreased with increasing PEG fraction, indicating that PEG competes with and displaces FITC-DEX from the membrane surface. These results suggest that PEG has a higher membrane affinity than DEX. Error bars represent standard deviations from two independent experiments using two separate SUV preparations.

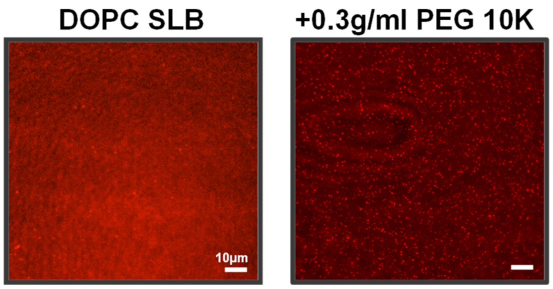

**Figure S7:** Representative TIRF images of supported lipid bilayers composed of 99.9 mol% DOPC+ 0.1 mol% RhoB-PE before polymer addition (left) and after addition of PEG (10 kDa, 0.3 g/mL) (right). PEG 10K did not induce detectable membrane patch formation; only small puncta were observed throughout the membrane. Scale bars, 10 μm.

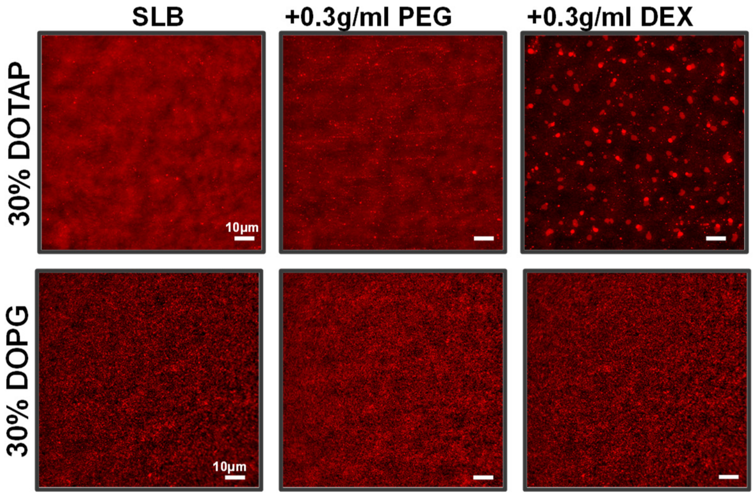

**Figure S8:** Representative TIRF images of supported lipid bilayers composed of 70 mol% DOPC + 30 mol% DOTAP + 0.1 mol% RhoB-PE (top row) or 70 mol% DOPC + 30 mol% DOPG + 0.1 mol% RhoB-PE (bottom row) before polymer addition (left), after addition of PEG (0.3 g/mL) (center), and after addition of DEX (0.3 g/mL) (right). DEX induced extensive micron-sized patch formation on DOTAP-containing membranes, whereas PEG produced no detectable membrane reorganization. In contrast, neither PEG nor DEX induced patch formation on DOPG-containing membranes. Scale bars, 10 μm.

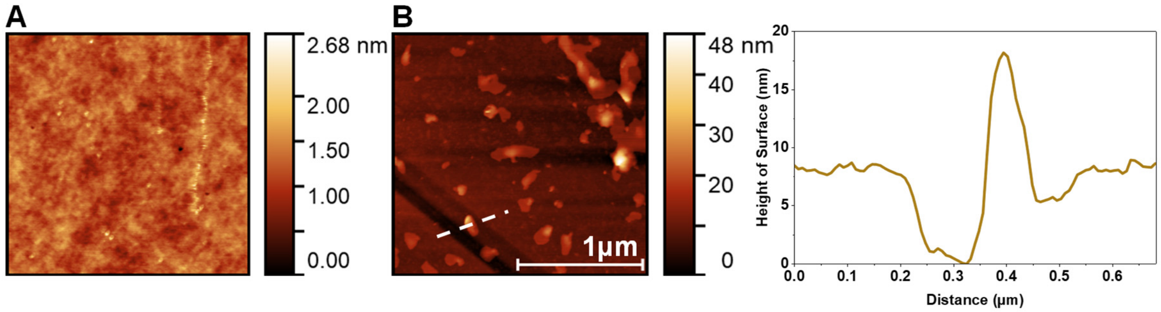

**Figure S9**: AFM characterization of DEX-induced membrane patches on supported lipid bilayers composed of 99.9 mol% DOPC and 0.1 mol% RhoB-PE. (A) Representative AFM image of the supported lipid bilayer before polymer addition, showing a laterally homogeneous membrane. (B) Representative AFM image after addition of 0.3 g/mL DEX, showing the formation of patches on the bilayer. The dashed line indicates the position of the height profile shown on the right. The intentionally introduced scratch exposes the underlying glass substrate, allowing measurement of both the supported lipid bilayer thickness and the additional height of the DEX-induced membrane patch. Scale bar, 1 μm.

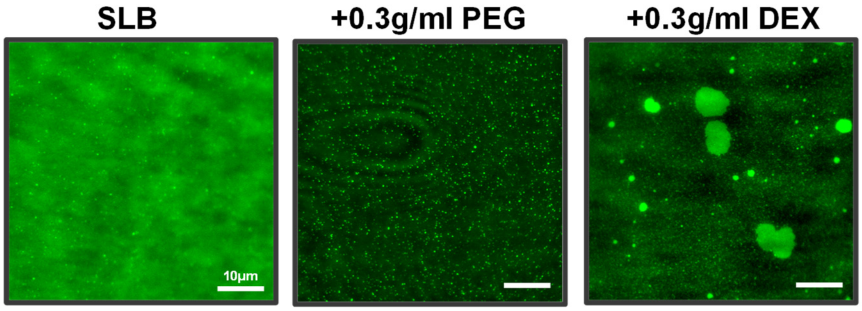

**Figure S10**: Representative TIRF images of a supported lipid bilayer composed of 99.5 mol% DOPC and 0.5 mol% BODIPY PC before polymer addition (left), after addition of PEG (0.3 g/mL) (center), and after addition of DEX (0.3 g/mL) (right). Similar DEX-induced patch formation was observed with BODIPY PC, demonstrating that the observed membrane reorganization is independent of the fluorophore and its membrane location. PEG did not induce patch formation, although small punctas were occasionally observed throughout the membrane. Scale bars, 10 μm.

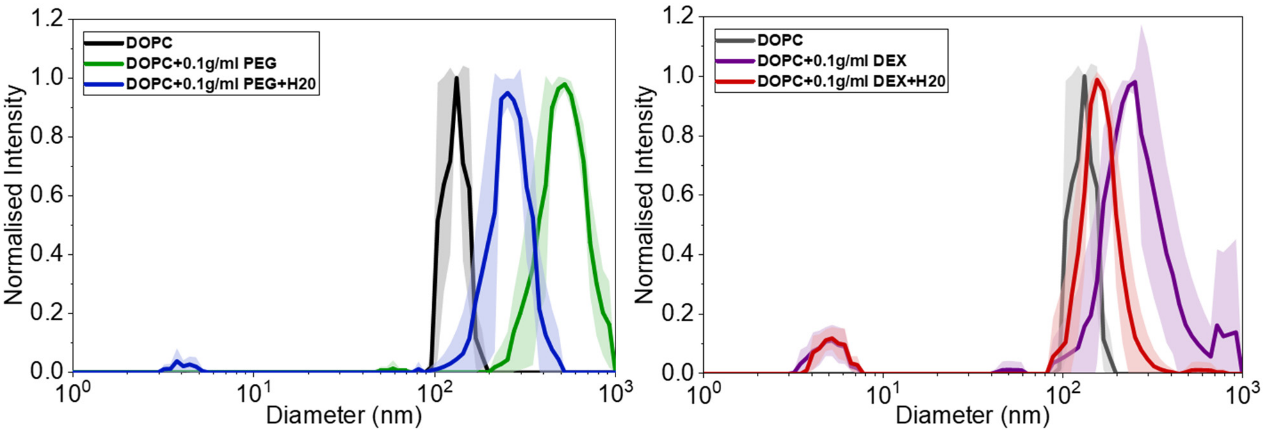

**Figure S11:** DLS size distributions of SUVs composed of 99.5 mol% DOPC and 0.5 mol% Laurdan in the absence and presence of PEG (0.1 g/mL) (left) or DEX (0.1 g/mL) (right). Addition of either polymer increased the apparent hydrodynamic diameter of the SUVs, indicating polymer-induced vesicle clustering. Following a subsequent 1:1 dilution with water, the average vesicle size decreased toward that of the untreated SUVs, indicating that the clustering was largely reversible. The increase in apparent vesicle size was more pronounced for PEG than for DEX, consistent with the stronger depletion-induced clustering reported for PEG-containing solutions. Error bars represent standard deviations from three independent experiments using two separate SUV preparations.

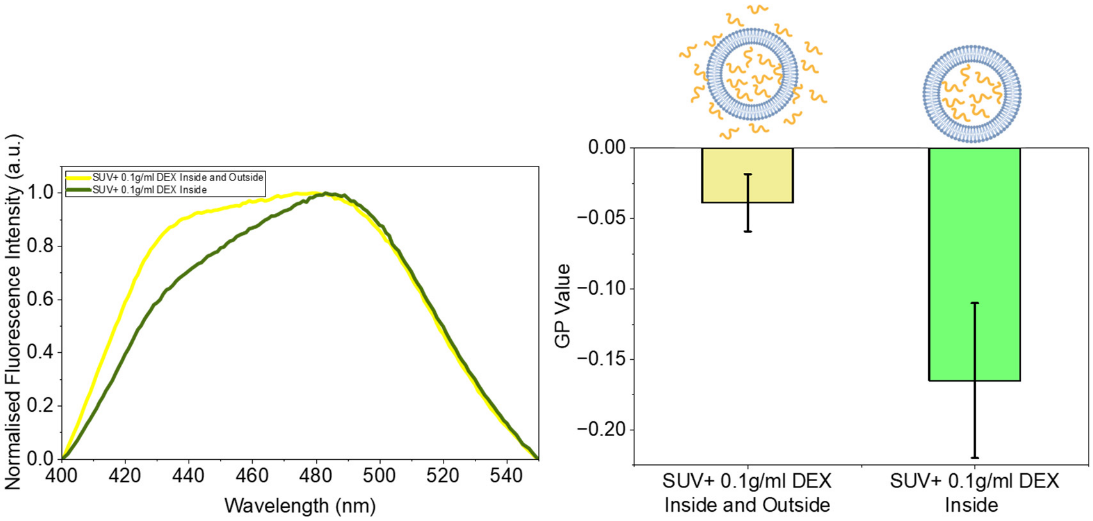

**Figure S12:** Laurdan emission spectra and corresponding GP values of SUVs composed of 99.5 mol% DOPC and 0.5 mol% Laurdan containing 0.1 g/mL DEX encapsulated within the vesicle lumen. Addition of 0.1 g/mL DEX to the external solution produced a spectral shift and GP increase comparable to those observed for water-filled SUVs upon external DEX addition (Figure 3C). These results indicate that the polymer-induced changes in Laurdan fluorescence are not primarily driven by an osmotic pressure difference across the vesicle membrane. Error bars represent standard deviations from three independent experiments using two separate SUV preparations.

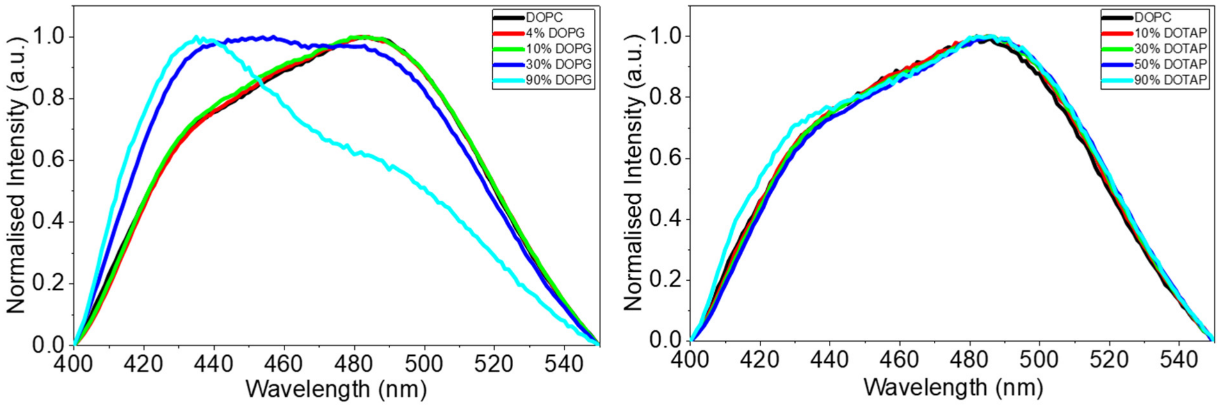

**Figure S13:** Laurdan emission spectra of SUVs composed of DOPC with increasing amounts of DOPG (0, 4, 10, 30, and 90 mol%; left) or DOTAP (0, 10, 30, 50, and 90 mol%; right), with all SUVs containing 0.5 mol% Laurdan, prior to DEX addition. The emission spectra were broadly similar within each lipid series, providing a baseline for comparison with the DEX-induced spectral changes shown in Figure 3 of the main text. The exception was SUVs containing 30 mol% and 90 mol% DOPG, which exhibited noticeable spectral differences prior to polymer addition. Previous studies have shown that PG headgroups can participate in extensive hydrogen-bonding interactions through their glycerol moieties, including both PG–PG and PG–water hydrogen bonds.[11, 12] These interactions become increasingly important at higher PG concentrations and can create a more ordered environment. As a result, water molecules surrounding Laurdan experienced reduced dipolar relaxation, leading to a more pronounced, blue-shifted emission near 440 nm. The absence of a similar effect at 4 mol% and 10 mol% DOPG suggests that a threshold PG concentration may be required before these collective hydration effects become significant[13].

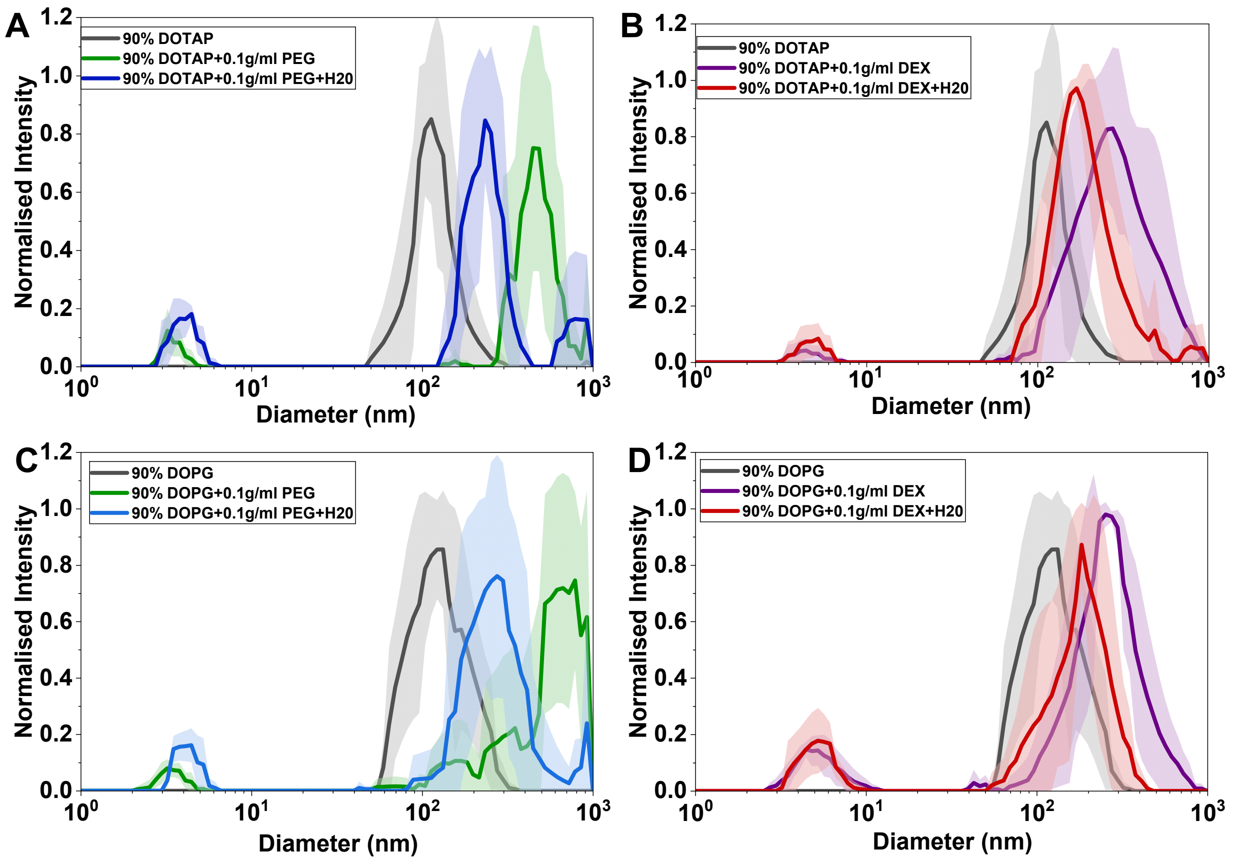

**Figure S14:** DLS size distributions of SUVs composed of 9.5 mol% DOPC + 90 mol% DOTAP + 0.5 mol% Laurdan (A, B) and 9.5 mol% DOPC + 90 mol% DOPG + 0.5 mol% Laurdan (C, D) in the presence of PEG (0.1 g/mL) (A, C) or DEX (0.1 g/mL) (B, D). Addition of either polymer increased the apparent hydrodynamic diameter of the SUVs, consistent with polymer-induced vesicle clustering. Following a subsequent 1:1 dilution with water, the average vesicle size decreased toward that of the untreated SUVs, indicating that the clustering process was largely reversible. Similar behavior was observed for both DOTAP- and DOPG-rich SUVs, suggesting that polymer-induced vesicle clustering is comparable across membrane compositions and does not account for the composition-dependent membrane responses observed in the main text.

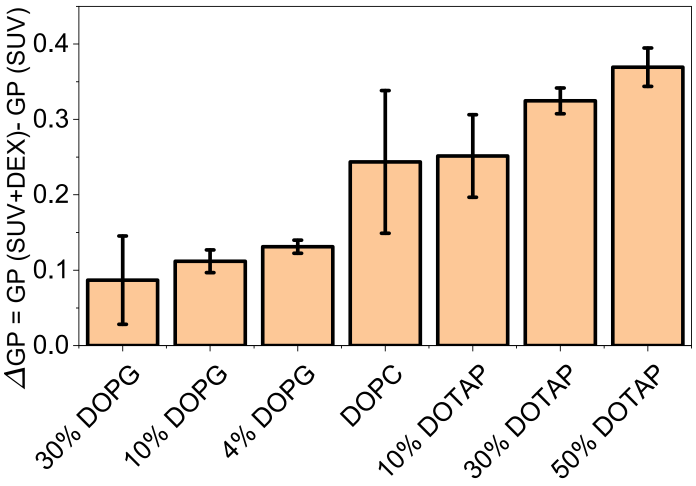

**Figure S15:** ΔGP values of SUVs composed of DOPC, DOPC/DOPG (4, 10, and 30 mol% DOPG), and DOPC/DOTAP (10, 30, 50, and 90 mol% DOTAP), with all SUVs containing 0.5 mol% Laurdan, measured in the presence of 0.1 g/mL DEX and 500 mM NaCl. A trend like that observed in water was retained under high-salt conditions, with larger ΔGP values for DOTAP-containing membranes. These results indicate that the membrane composition-dependent GP changes upon DEX intearctions are not primarily driven by electrostatic interactions between the polymer and the membrane. Error bars represent standard deviations from three independent experiments using two separate SUV preparations.

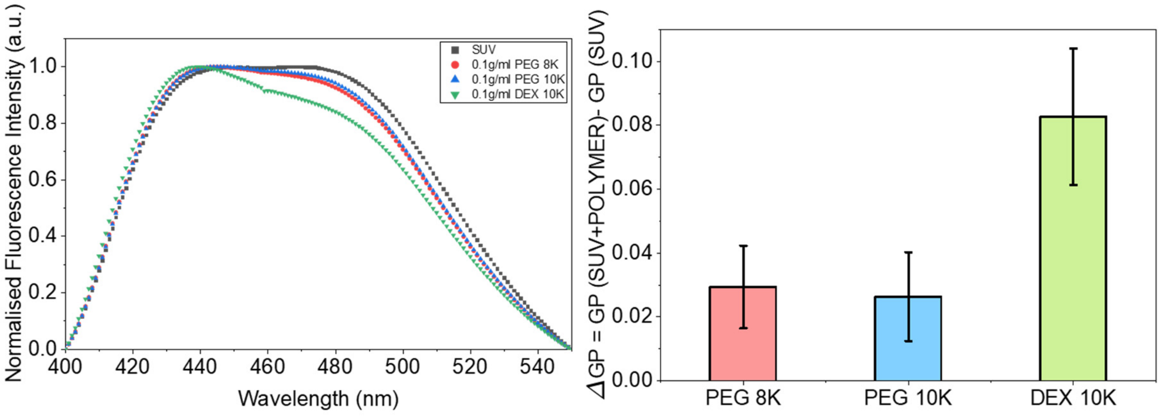

**Figure S16.** Comparison of the effects of PEG 8 kDa, PEG 10 kDa, and DEX 10 kDa on Laurdan emission spectra and GP values. Laurdan emission spectra of SUVs composed of 99.5 mol% DOPC and 0.5 mol% Laurdan before polymer addition and after addition of 0.1 g/mL PEG 8 kDa, PEG 10 kDa, or DEX 10 kDa (left). Corresponding changes in GP values (ΔGP) are shown on the right. PEG 8 kDa and PEG 10 kDa produced similar spectral shifts and comparable ΔGP values, whereas DEX 10 kDa induced a substantially larger GP change. Error bars represent standard deviations from two independent experiments using two separate SUV preparations.

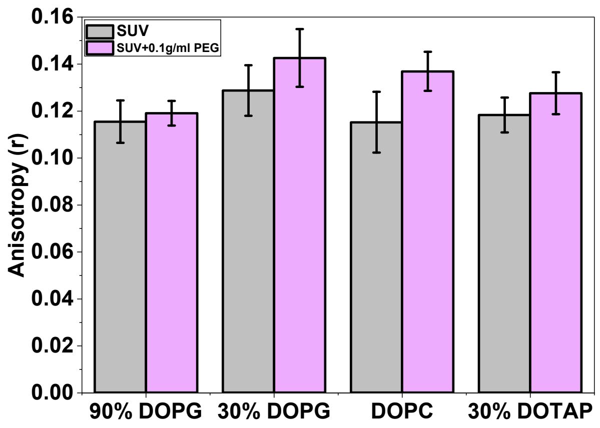

**Figure S17.** Fluorescence anisotropy of SUVs composed of DOPC, DOPC/DOPG (30 and 90 mol% DOPG), and DOPC/DOTAP (30 mol% DOTAP), with all SUVs containing 0.5 mol% DPH, measured before and after addition of 0.1 g/mL PEG. PEG produced only small changes in DPH anisotropy across all membrane compositions, indicating minimal changes in hydrocarbon chain packing. Error bars represent standard deviations from three independent experiments using two separate SUV preparations.

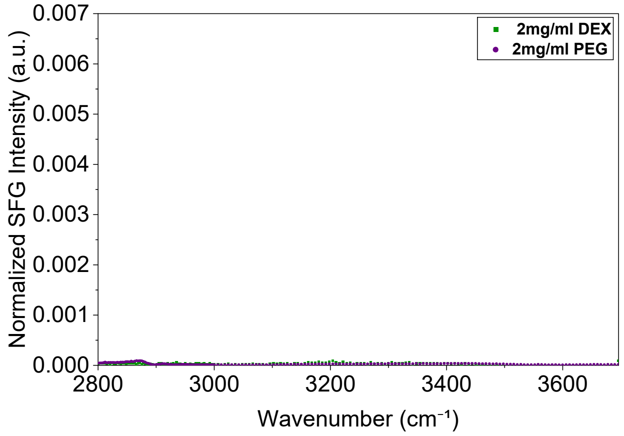

**Figure S18**: VSFG spectra of 2 mg/mL DEX and 2 mg/mL PEG in the aqueous solution in the absence of a lipid monolayer. Neither polymer produced a detectable SFG signal.

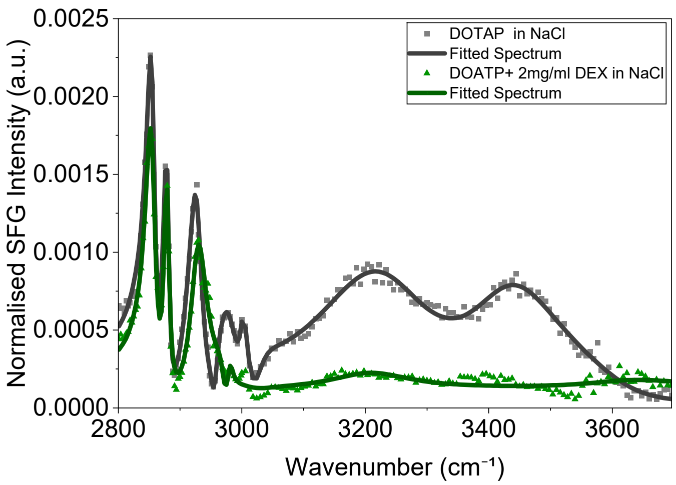

**Figure S19:** VSFG spectra of a 100 mol% DOTAP monolayer before and after addition of 2 mg/mL DEX in the presence of 150 mM NaCl in the aqueous subphase. The suppression of the interfacial O–H stretching bands persists under high-salt conditions, indicating that the DEX-induced changes in interfacial hydration are not driven by ionic effects. Solid lines represent the fitted spectra.

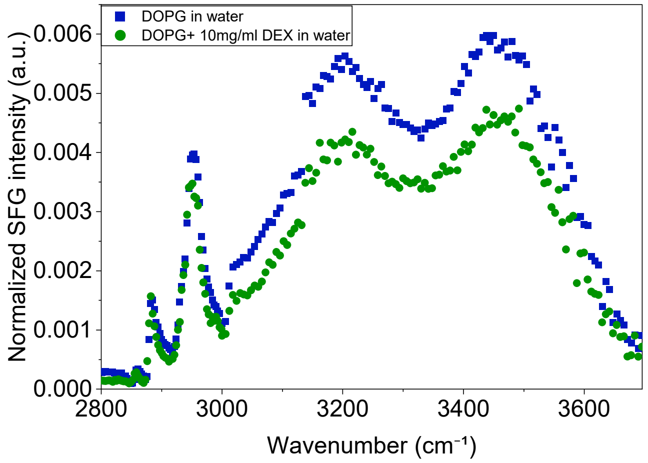

**Figure S20:** VSFG spectra of a 100 mol% DOPG monolayer before and after addition of 10 mg/mL DEX in the aqueous subphase. The decrease in the O–H stretching intensity at high DEX concentration indicates a modest perturbation of the interfacial hydration layer.

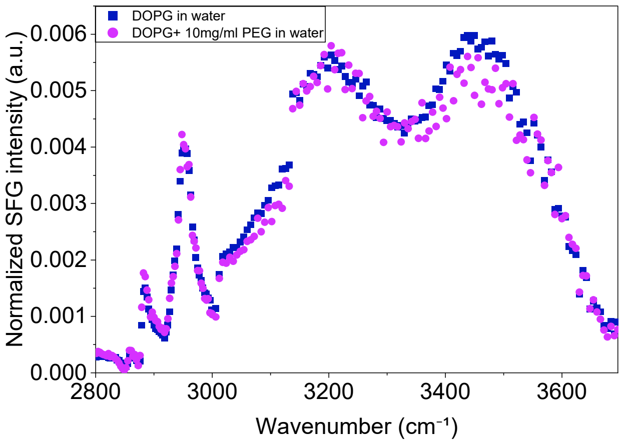

**Figure S21:** VSFG spectra of a 100 mol% DOPG monolayer before and after addition of 10 mg/mL PEG in the aqueous subphase. The slight increase in the O–H stretching intensity is consistent with a modest increase in interfacial hydration upon PEG adsorption.

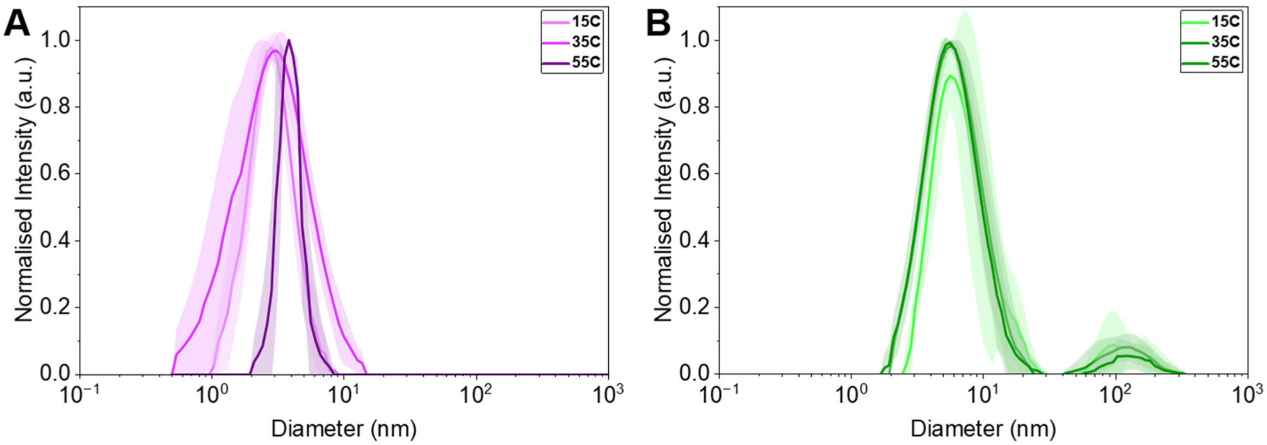

**Figure S22:** Hydrodynamic diameter distributions of PEG (A) and DEX (B) measured by dynamic light scattering (DLS) at 5, 35, and 65 °C. Only minor changes in the hydrodynamic diameter distributions of either polymer were observed over this temperature range, indicating negligible changes in polymer self-association prior to membrane binding. Shaded regions represent the standard deviations from three independent measurements.

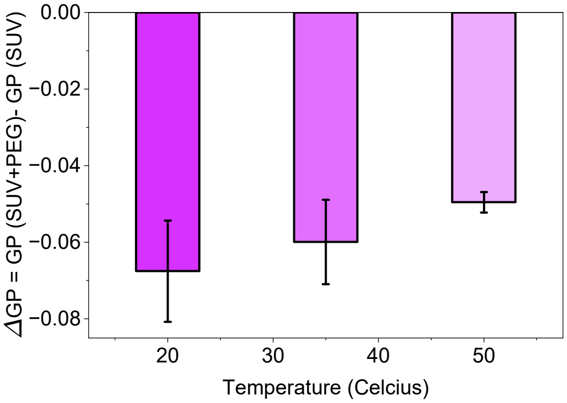

**Figure S23:** Temperature dependence of the PEG-induced change in Laurdan generalized polarization (GP) for SUVs composed of 69.5 mol% DOPC+30 mol% DOPG +0.5 mol% Laurdan following addition of 0.1 g/mL PEG. While the PEG-induced decrease in GP became smaller with increasing temperature, the amount of membrane-bound PEG remained consistent across the temperature (Figure 6). This result indicates that, at higher temperature, there can be an entropy gain from weaker hydrogen bond network at the PEG-PG interface. This increase in entropy can offset the weakening of water-mediated hydrogen bonding, resulting in little overall change in the binding free energy and, therefore, minimal temperature dependence of PEG binding. Error bars represent standard deviations from two independent experiments using two separate SUV preparations.

**Table S1:** Hydrodynamic diameter, zeta potential, and solution conductivity of fluorescently labeled and unlabeled polymers used in this study. Measurements were performed in ultrapure water (18.2 MΩ·cm at 25 °C) at the concentrations used for imaging experiments. All polymers exhibited zeta potentials within the approximately neutral range (−5 to +5 mV)[14]. The fluorescently labeled polymers displayed hydrodynamic diameters, zeta potential, and solution conductivities comparable to those of their corresponding unlabeled polymers, indicating that fluorophore labeling did not substantially alter the physicochemical properties of the polymers.

| Concentration (g/ml) | Polymer | Average Diameter (nm) | SD | Zeta Potential (mV) | SD | Solution Conductivity (mS/cm) | SD |
|---|---|---|---|---|---|---|---|
| 0.1 | PEG 8K | 3.28 | 0.19 | -1.76 | 0.92 | 0.03 | 0.0004 |
| | DEX 10K* | 42.27 (With aggregates) | 52.43 | -3.27 | 0.63 | 0.04 | 0.0001 |
| | | 7.29 (Without aggregates) | 1.19 | | | | |
| 0.001 | PEG 8K | 4.76 | 1.06 | -0.99 | 1.97 | 0.006 | 0.0003 |
| | DEX 10K | 6.96 | 5.86 | 0.46 | 1.25 | 0.007 | 0.0007 |
| | RhoB-PEG | 2.38 | 1.15 | -2.03 | 0.94 | 0.017 | 0.0005 |
| | FITC-DEX | 5.07 | 0.83 | -0.2 | 0.41 | 0.011 | 0.0006 |

* DEX 10K exhibited a secondary population with a larger apparent hydrodynamic diameter, likely due to intra- and intermolecular hydrogen-bonding interactions involving the hydroxyl groups of dextran. These interactions can promote the formation of a small population of aggregates in solution. The dominant particle population remained centered at approximately 7.3 nm.

**Table S2:** Peak positions, amplitudes ($A_n$), and linewidths ($\Gamma_n$) obtained from fitting the SFG spectra of 100 mol% DOTAP monolayer before and after addition of 2 mg/mL DEX. Peak assignments are based on the vibrational modes of the lipid C–H stretching region and the interfacial O–H stretching region. Values are reported as mean ± standard deviation from two independent measurements.

| Wavenumber ($cm^{-1}$) | DOTAP before DEX<br>$A_n$<br>$\Gamma_n$ ($cm^{-1}$) | Wavenumber ($cm^{-1}$) | DOTAP + 2 mg/mL DEX<br>$A_n$<br>$\Gamma_n$ ($cm^{-1}$) |
|---|---|---|---|
| 2857 ± 1<br>($CH_2$ sym.) | -0.23 ± 0.01<br>7.6 ± 0.5 | 2857 ± 1.5<br>($CH_2$ sym.) | -0.23 ± 0.03<br>8.6 ± 0.4 |
| 2880 ± 0.6<br>($CH_3$ sym.) | -0.20 ± 0.03<br>6.0 ± 0.5 | 2880 ± 0.6<br>($CH_3$ sym.) | -0.16 ± 0.02<br>5.5 ± 0.1 |
| 2927 ± 0.5<br>($CH_2$-FR) | -0.47 ± 0.08<br>13.4 ± 0.3 | 2929 ± 1.2<br>($CH_2$-FR) | -0.33 ± 0.21<br>12.1 ± 3.4 |
| 2941 ± 1.2<br>($CH_3$-FR) | -0.024 ± 0.02<br>3.4 ± 2 | 2944 ± 1.2<br>($CH_3$-FR) | -0.036 ± 0.01<br>4.4 ± 5.2 |
| 2959 ± 0.6<br>($CH_3$ asym.) | -0.1 ± 0<br>6.6 ± 0.2 | 2957 ± 2.5<br>($CH_3$ asym.) | 0.028 ± 0.02<br>5.3 ± 9.3 |
| 2983 ± 1<br>(Choline $CH_3$ sym.) | -0.023 ± 0.01<br>5.3 ± 0.9 | 2981 ± 4.4<br>(Choline $CH_3$ sym.) | -0.09 ± 0.06<br>13.8 ± 5.4 |
| 3012 ± 3.2<br>(vinyl CH) | -0.57 ± 0.4<br>29.6 ± 13.1 | 3009 ± 1<br>(vinyl CH) | -0.22 ± 0.09<br>15.4 ± 2.2 |
| 3025 ± 0<br>(Choline $CH_3$ asym.) | 0.12 ± 0.06<br>12.3 ± 1.9 | 3021 ± 6.1<br>(Choline $CH_3$ asym.) | 0.18 ± 0.28<br>16.1 ± 2.5 |
| 3220 ± 19.8<br>(OH 1) | -7.6 ± 0.62<br>157 ± 16 | 3213 ± 37<br>(OH 1) | -3.2 ± 0.91<br>175.9 ± 32.2 |
| 3433 ± 9.3<br>(OH 2) | -3.7 ± 0.84<br>100.2 ± 10.8 | 3436 ± 18.1<br>(OH 2) | -0.7 ± 1.56<br>63.3 ± 30.4 |
| 3604 ± 4.2<br>(OH 3) | 2.8 ± 0.45<br>89.5 ± 4.5 | 3619 ± 9.2<br>(OH 3) | 0.9 ± 0.21<br>63.3 ± 29.3 |

**Table S3:** Peak positions, amplitudes ($A_n$), and linewidths ($\Gamma_n$) obtained from fitting the SFG spectra of 100 mol% DOTAP monolayer before and after addition of 2 mg/mL PEG. Peak assignments are based on the vibrational modes of the lipid C–H stretching region and the interfacial O–H stretching region. Values are reported as mean ± standard deviation from two independent measurements.

| Wavenumber ($cm^{-1}$) | DOTAP before PEG<br>$A_n$<br>$\Gamma_n$ ($cm^{-1}$) | Wavenumber ($cm^{-1}$) | DOTAP + 2 mg/mL PEG<br>$A_n$<br>$\Gamma_n$ ($cm^{-1}$) |
|---|---|---|---|
| 2857 ± 0<br>($CH_2$ sym.) | -0.23 ± 0.01<br>7.6 ± 0.5 | 2857 ± 0<br>($CH_2$ sym.) | -0.25 ± 0.0<br>7.7 ± 0.5 |
| 2880 ± 0<br>($CH_3$ sym.) | -0.20 ± 0.03<br>6.0 ± 0.5 | 2880 ± 0<br>($CH_3$ sym.) | -0.23 ± 0.01<br>6.1 ± 0.4 |
| 2928 ± 0.7<br>($CH_2$-FR) | -0.47 ± 0.08<br>13.4 ± 0.3 | 2928 ± 0.7<br>($CH_2$-FR) | -0.55 ± 0.1<br>13.8 ± 1 |
| 2942 ± 1.4<br>($CH_3$-FR) | -0.024 ± 0.02<br>3.4 ± 2 | 2942 ± 0<br>($CH_3$-FR) | -0.03 ± 0.01<br>3.5 ± 0.5 |
| 2959 ± 0.7<br>($CH_3$ asym.) | -0.1 ± 0<br>6.6 ± 0.2 | 2957 ± 0.7<br>($CH_3$ asym.) | 0.14 ± 0.01<br>7.8 ± 0.4 |
| 2983 ± 1.4<br>(Choline $CH_3$ sym.) | -0.023 ± 0.01<br>5.3 ± 0.9 | 2982 ± 0.7<br>(Choline $CH_3$ sym.) | -0.04 ± 0.02<br>7.6 ± 1.7 |
| 3010 ± 3.5<br>(vinyl CH) | -0.57 ± 0.4<br>29.6 ± 13.1 | 3012 ± 4.2<br>(vinyl CH) | -0.61 ± 0.51<br>20 ± 5.5 |
| 3025 ± 0<br>(Choline $CH_3$ asym.) | 0.12 ± 0.06<br>12.3 ± 1.9 | 3017 ± 6.4<br>(Choline $CH_3$ asym.) | 0.19 ± 0.25<br>10.1 ± 0.1 |
| 3229 ± 15.6<br>(OH 1) | -7.6 ± 0.62<br>157 ± 16 | 3211 ± 7.1<br>(OH 1) | -8.1 ± 1.8<br>121.1 ± 18.7 |
| 3437 ± 12.7<br>(OH 2) | -3.7 ± 0.84<br>100.2 ± 10.8 | 3429 ± 4.9<br>(OH 2) | -6 ± 0.07<br>122.9 ± 0.8 |
| 3602 ± 2.1<br>(OH 3) | 2.8 ± 0.45<br>89.5 ± 4.5 | 3606 ± 9.2<br>(OH 3) | 3.4 ± 0.14<br>84.3 ± 7.5 |

**Table S4:** Peak positions, amplitudes ($A_n$), and linewidths ($\Gamma_n$) obtained from fitting the SFG spectra of 100 mol% DOPG monolayer before and after addition of 2 mg/mL DEX. Peak assignments are based on the vibrational modes of the lipid C–H stretching region and the interfacial O–H stretching region. Values are reported as mean ± standard deviation from two independent measurements.

| Wavenumber ($cm^{-1}$) | DOPG before DEX<br>$A_n$<br>$\Gamma_n$ ($cm^{-1}$) | Wavenumber ($cm^{-1}$) | DOPG + 2 mg/mL DEX<br>$A_n$<br>$\Gamma_n$ ($cm^{-1}$) |
|---|---|---|---|
| 2856 ± 0.9<br>($CH_2$ sym.) | -0.08 ± 0.03<br>7.2 ± 2.2 | 2857 ± 0.7<br>($CH_2$ sym.) | -0.08 ± 0.03<br>5.6 ± 1.4 |
| 2880 ± 1.1<br>($CH_3$ sym.) | -0.15 ± 0.03<br>5.3 ± 0.2 | 2880 ± 0<br>($CH_3$ sym.) | -0.19 ± 0.04<br>6.7 ± 0.3 |
| 2928 ± 2<br>($CH_2$-FR) | -0.32 ± 0.9<br>14.8 ± 2 | 2930 ± 0<br>($CH_2$-FR) | -0.48 ± 0<br>17.6 ± 3.7 |
| 2944 ± 0.8<br>($CH_3$-FR) | -0.22 ± 0.26<br>10.3 ± 4.3 | 2944 ± 0.7<br>($CH_3$-FR) | -0.14 ± 0.16<br>8.0 ± 2.7 |
| 2959 ± 2.4<br>($CH_3$ asym.) | 0.58 ± 0.2<br>20.5 ± 2.5 | 2961 ± 0<br>($CH_3$ asym.) | 0.51 ± 0.12<br>20.2 ± 4.1 |
| 3007 ± 1.3<br>(vinyl CH) | -0.06 ± 0.01<br>5.7 ± 1 | 3009 ± 1.4<br>(vinyl CH) | -0.07 ± 0.01<br>5.8 ± 2.2 |
| 3208 ± 4.9<br>(OH 1) | 7.8 ± 0.78<br>127.4 ± 9.8 | 3212 ± 4.9<br>(OH 1) | 7.8 ± 0.5<br>123.4 ± 4.1 |
| 3423 ± 4<br>(OH 2) | 6.5 ± 0.9<br>125 ± 13.3 | 3423 ± 8.5<br>(OH 2) | 6.5 ± 0<br>118.6 ± 13.4 |
| 3621 ± 10.7<br>(OH 3) | -2.2 ± 0.83<br>90.6 ± 18.7 | 3613 ± 9.2<br>(OH 3) | -2.5 ± 0.14<br>96.3 ± 0.4 |

**Table S5:** Peak positions, amplitudes ($A_n$), and linewidths ($\Gamma_n$) obtained from fitting the SFG spectra of 100 mol% DOPG monolayer before and after addition of 2 mg/mL PEG. Peak assignments are based on the vibrational modes of the lipid C–H stretching region and the interfacial O–H stretching region. Values are reported as mean ± standard deviation from two independent measurements.

| Wavenumber ($cm^{-1}$) | DOPG before PEG<br>$A_n$<br>$\Gamma_n$ ($cm^{-1}$) | Wavenumber ($cm^{-1}$) | DOPG + 2 mg/mL PEG<br>$A_n$<br>$\Gamma_n$ ($cm^{-1}$) |
|---|---|---|---|
| 2856 ± 0.9<br>($CH_2$ sym.) | -0.08 ± 0.03<br>7.2 ± 2.2 | 2856 ± 0.7<br>($CH_2$ sym.) | -0.1 ± 0<br>6.1 ± 0.3 |
| 2880 ± 1.1<br>($CH_3$ sym.) | -0.15 ± 0.03<br>5.3 ± 0.2 | 2880 ± 0<br>($CH_3$ sym.) | -0.19 ± 0.01<br>5.4 ± 0.3 |
| 2928 ± 2<br>($CH_2$-FR) | -0.32 ± 0.9<br>14.8 ± 2 | 2927 ± 0<br>($CH_2$-FR) | -0.32 ± 0.01<br>14.2 ± 0.5 |
| 2944 ± 0.8<br>($CH_3$-FR) | -0.22 ± 0.26<br>10.3 ± 4.3 | 2943 ± 0<br>($CH_3$-FR) | -0.19 ± 0.02<br>7.5 ± 0.5 |
| 2959 ± 2.4<br>($CH_3$ asym.) | 0.58 ± 0.2<br>20.5 ± 2.5 | 2958 ± 0.7<br>($CH_3$ asym.) | 0.68 ± 0.04<br>22.2 ± 0.3 |
| 3007 ± 1.3<br>(vinyl CH) | -0.06 ± 0.01<br>5.7 ± 1 | 3007 ± 0.7<br>(vinyl CH) | -0.06 ± 0<br>5.1 ± 0.1 |
| 3208 ± 4.9<br>(OH 1) | 7.8 ± 0.78<br>127.4 ± 9.8 | 3207 ± 1.4<br>(OH 1) | 7.93 ± 0.2<br>124.2 ± 0 |
| 3423 ± 4<br>(OH 2) | 6.5 ± 0.9<br>125 ± 13.3 | 3422 ± 0.7<br>(OH 2) | 7 ± 0<br>143.9 ± 3.6 |
| 3621 ± 10.7<br>(OH 3) | -2.2 ± 0.83<br>90.6 ± 18.7 | 3618 ± 0<br>(OH 3) | -3 ± 0<br>94.8 ± 1.6 |

**Discussion:**

**Selection of PEG 8 kDa over PEG 10 kDa for PEG–DEX 10 kDa Aqueous Two-Phase Systems**

Polyethylene glycol with a molecular weight of 8 kDa (PEG) and dextran with a molecular weight of 10 kDa (DEX) are among the most used polymer pairs in PEG–DEX aqueous two-phase systems and artificial cell studies[15]. PEG molecular weight is known to influence the phase behavior of PEG–DEX aqueous two-phase systems. Increasing polymer molecular weight enhances the phase-forming ability of the system[16]. In addition, higher molecular weight PEGs generally exhibit higher solution viscosities[17]. We compared PEG 8K and PEG 10K in our system and found that they exhibited similar hydrodynamic diameters by DLS (3.29 ± 0.24 nm for PEG 8 kDa and 3.49 ± 0.11 nm for PEG 10 kDa). They also produced similar changes in Laurdan emission spectra and GP values (Figure S15). Therefore, PEG 8 kDa was selected for the remainder of this study. PEG 8K solutions were also easier to prepare and handle because of their lower viscosity.

**Rationale for Selecting FITC–DEX and RhoB–PEG as Fluorescent Polymer Probes**

FITC-DEX and RhoB-PEG were used as fluorescent tracers for DEX and PEG, respectively. FITC-DEX has been widely used in permeability, diffusion, and transport studies because its low degree of labeling (0.003–0.02 mol FITC per mol glucose residue) introduces only minimal changes to the properties of the parent DEX. As a result, FITC-DEX is expected to retain the behavior of unlabeled DEX while enabling fluorescence imaging.[18, 19] Furthermore, FITC is relatively hydrophilic and, at the low degree of labeling used here, minimally perturbs the highly hydrated nature of DEX. In contrast, the more hydrophobic RhoB label can introduce additional hydrophobic interactions. Because PEG possesses inherent amphiphilic character,[20] the RhoB label is expected to have a smaller relative effect on PEG than on DEX. To evaluate the suitability of fluorescent dextran tracers, we compared the partitioning behavior of FITC-DEX and Rhodamine B-labelled Dextran (RhoB-DEX) in a PEG/DEX aqueous two-phase system consisting of PEG (0.5 g/mL) and DEX (0.5 g/mL). Following phase separation and centrifugation, FITC-DEX partitioned almost exclusively into the DEX-rich phase, whereas RhoB-DEX exhibited detectable partitioning into both the DEX-rich and PEG-rich phases (Figure S22. In addition, detailed structural information regarding the degree of substitution and branching characteristics was not available for the RhoB-DEX product used in this study. Therefore, FITC-DEX was selected as the fluorescent DEX analogue throughout this work. RhoB-PEG was selected as a commercially available fluorescent PEG derivative that has been widely used for fluorescence imaging and tracking of PEG-containing systems.

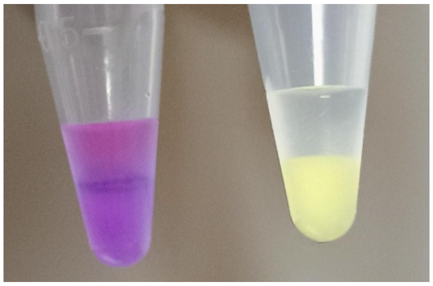

**Figure S22:** Partitioning behavior of fluorescently labeled dextran polymers in a PEG–DEX aqueous two-phase system. FITC-DEX (right) partitioned almost exclusively into the DEX-rich phase, whereas RhoB-DEX (left) exhibited detectable partitioning into both the DEX-rich and PEG-rich phases. These results demonstrate that the fluorophore can influence dextran partitioning behavior in PEG–DEX phase-separated systems.

**Sensitivity of Laurdan Fluorescence to Interfacial Hydration and Lipid Packing**

Laurdan (6-dodecanoyl-2-dimethylaminonaphthalene) is an amphiphilic fluorescent probe originally developed by Weber to study solvent dipolar relaxation. Since then, it has become one of the most widely used probes for characterizing membrane hydration and lipid organization.[4, 7] Previous experimental and computational studies have shown that Laurdan localizes near the glycerol backbone region of phospholipid membranes, with the fluorophore positioned approximately 1.0–1.2 nm below the bilayer interface. The dimethyl amino group of Laurdan resides approximately in the same plane as the lipid carbonyl groups, placing the probe at the boundary between the hydrophobic membrane core and the interfacial hydration layer.[21] This location is particularly important because it allows Laurdan to simultaneously sense both membrane packing and the interfacial hydration environment surrounding the lipid carbonyl and headgroup region.

The interfacial hydration layer differs substantially from bulk water. Water molecules in the lipid headgroup region interact with the choline, phosphate, and carbonyl groups of the membrane and participate in an extended hydrogen-bonding network. Water molecules located deeper within the headgroup region exhibit reduced mobility and increased stability relative to bulk water.[22, 23] Since Laurdan resides within this interfacial region, its fluorescence is highly sensitive to changes in the local hydration environment and water dynamics at the membrane interface.

The spectral sensitivity of Laurdan originates from an intramolecular charge-transfer process that occurs upon excitation. Laurdan contains an electron-donating dimethyl amino group and an electron-withdrawing carbonyl group. When the fluorophore is excited from the ground state ($S_0$) to the excited state ($S_1$), electron density redistributes within the molecule, resulting in a substantially larger excited-state dipole moment than in the ground state. This newly generated dipole perturbs the surrounding medium and gives rise to the solvatochromic behavior of Laurdan.[7]

In a nonpolar environment, where few surrounding dipoles are available to respond to the excited fluorophore, the excited state remains largely unstabilized. Under these conditions, Laurdan emits directly from the excited state ($S_1$). In a polar environment, however, surrounding dipoles can respond to the excited-state dipoles. According to Onsager reaction field theory, the excited Laurdan molecule polarizes the surrounding medium, and the surrounding dipoles subsequently generate a reaction field that stabilizes the excited state. This initial stabilization produces a lower-energy excited state (non-relaxed excited state). If emission occurs from this state before further dipolar reorientation takes place, Laurdan emits at approximately 440 nm.[7]

A second level of stabilization can occur if the surrounding dipoles are sufficiently mobile. Water molecules located near the fluorophore can physically reorient around the excited-state dipole during the fluorescence lifetime of Laurdan. This process, known as dipolar relaxation, further lowers the energy of the excited state and generates the relaxed excited state. Emission from this relaxed state occurs at a lower energy level and therefore appears at longer wavelengths, typically around 490 nm. Thus, polarity determines the initial stabilization of the excited state, whereas dipolar relaxation determines the extent of additional stabilization prior to emission.[7]

When Laurdan is incorporated into lipid membranes, both membrane packing and interfacial hydration influence its emission properties. In less fluid membranes, where the number of interfacial water molecules is lower and lipids are more tightly packed, the polarity of the medium and the extent of dipolar relaxation is reduced. Under these conditions, Laurdan exhibits a blue-shifted emission spectrum with increased intensity near 440 nm. In contrast, in more fluid membranes, where the number of interfacial water molecules is greater and lipids are more loosely packed, polarity of the medium increases and the dipolar relaxation become more efficient, leading to increased emission near 490 nm. Thus, Laurdan reports the combined effects of membrane hydration and lipid packing, which are often collectively described as membrane fluidity.[21] To quantitatively describe these spectral changes, Parasassi and co-workers introduced the term generalized polarization (GP), which compares the fluorescence intensities at 440 and 490 nm. An increase in GP corresponds to increased emissions near 440 nm (blue-shifted emission) and is generally associated with a less fluid membrane environment. Conversely, a decrease in GP corresponds to increased emissions near 490 nm (red-shifted emission) and is generally associated with a more fluid membrane environment.[4, 7]

In the present study, however, several observations suggest that the polymer-induced GP changes primarily originate from changes in interfacial hydration rather than changes in membrane packing. FRAP measurements revealed little change in lipid mobility following polymer addition, and DPH anisotropy measurements showed minimal changes in acyl-chain packing. If the observed GP increase arose primarily from changes in membrane packing, corresponding changes would be expected in these measurements. In contrast, SFG measurements revealed a substantial decrease in the interfacial O–H signal following polymer addition, indicating significant perturbation of membrane hydration at the membrane interface.

Taking together, these results suggest that the increase in Laurdan GP observed in this study is dominated by changes in interfacial hydration. Polymer binding reduces the number of water molecules surrounding Laurdan within the glycerol backbone region, thereby restricting dipolar relaxation of the excited fluorophore. The resulting reduction in excited-state stabilization produces the observed blue shift in Laurdan emission and corresponding increase in GP. Therefore, although Laurdan is often described as a reporter of membrane fluidity, our combined FRAP, DPH anisotropy, and SFG measurements indicate that Laurdan is primarily reporting changes in interfacial hydration under the conditions investigated here.

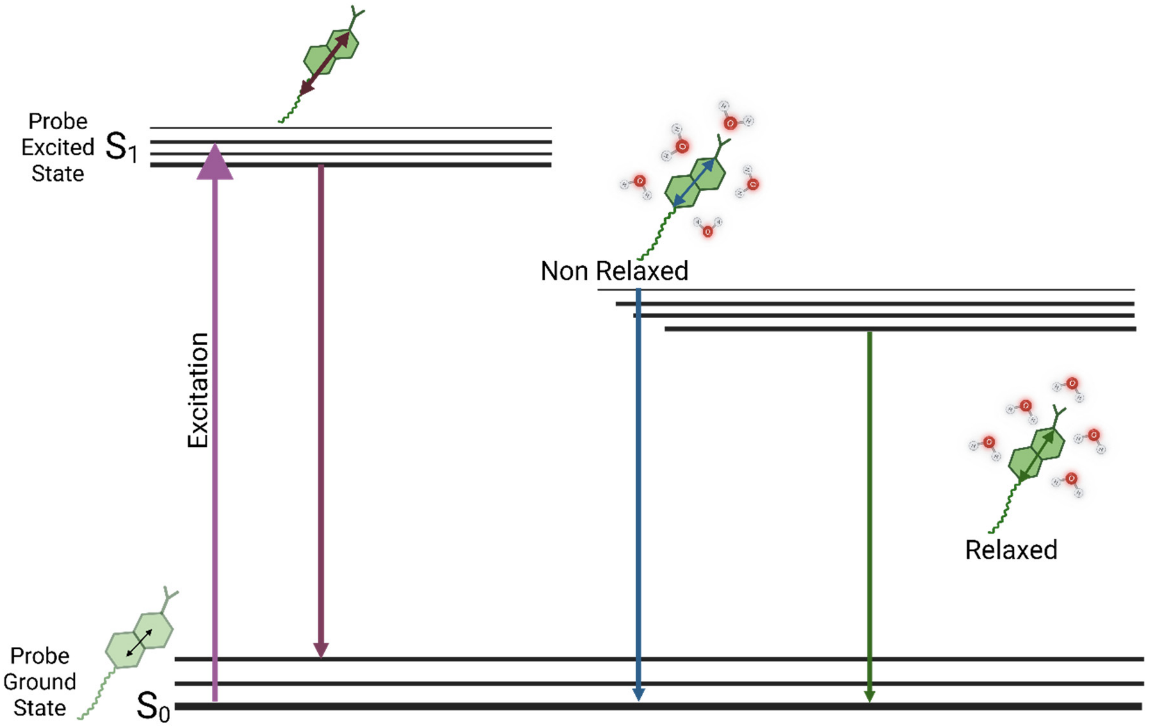

**Figure S23:** Schematic illustration of the photophysical mechanism of Laurdan. Upon excitation, Laurdan undergoes intramolecular charge transfer, generating a larger excited-state dipole. In a nonpolar environment, emissions occur directly from the excited state. In a polar environment, the excited state is initially stabilized by the Onsager reaction field, producing the non-relaxed excited state (~440 nm). Subsequent dipolar relaxation of surrounding water molecules further stabilizes the excited state, resulting in red-shifted emission (~490 nm).
